# A Highly Immunogenic Measles Virus-based Th1-biased COVID-19 Vaccine

**DOI:** 10.1101/2020.07.11.198291

**Authors:** Cindy Hörner, Christoph Schürmann, Arne Auste, Aileen Ebenig, Samada Muraleedharan, Maike Herrmann, Barbara Schnierle, Michael D. Mühlebach

## Abstract

The COVID-19 pandemic is caused by severe acute respiratory syndrome coronavirus-2 (SARS-CoV-2) and has spread world-wide with millions of cases and hundreds of thousands of deaths to date. The gravity of the situation mandates accelerated efforts to identify safe and effective vaccines. Here, we generated measles virus (MeV)-based vaccine candidates expressing the SARS-CoV-2 spike glycoprotein (S). Insertion of the full-length S protein gene in two different MeV genomic positions resulted in modulated S protein expression. The variant with lower S protein expression levels was genetically stable and induced high levels of effective Th1-biased antibody and T cell responses in mice after two immunizations. In addition to neutralizing IgG antibody responses in a protective range, multifunctional CD8^+^ and CD4^+^ T cell responses with S protein-specific killing activity were detected. These results are highly encouraging and support further development of MeV-based COVID-19 vaccines.

**Author Contributions:** CH performed research, analyzed data, and wrote the paper; CS performed research and analyzed data; AA performed research and analyzed data; AE performed research and analyzed data; SM performed research, analyzed data, and wrote the paper; MH developed the bioinformatics pipeline and analyzed data; BS contributed new reagents and concepts; MDM designed and supervised research, analyzed data and wrote the paper; all authors read, corrected and approved the final manuscript.

**Significance Statement:** The COVID-19 pandemic has caused hundreds of thousands of deaths, yet. Therefore, effective vaccine concepts are urgently needed. In search for such a concept, we have analysed a measles virus-based vaccine candidate targeting SARS-CoV-2. Using this well known, safe vaccine backbone, we demonstrate here induction of functional immune responses in both arms of adaptive immunity with the desired immune bias. Therefore, occurrence of immunopathologies such as antibody-dependent enhancement or enhanced respiratory disease is rather unlikely. Moreover, the candidate still induces immunity against the measles, recognized as a looming second menace, when countries are entrapped to stop routine vaccination campaigns in the face of COVID-19. Thus, a bivalent measles-based COVID-19 vaccine could be the solution for two significant public health threats.

## 76 Introduction

Severe acute respiratory syndrome coronavirus-2 (SARS-CoV-2) belongs to *Coronaviridae* family and emerged towards the end of 2019 as causative agent of pneumonia in the Hubei province in China (1). The World Health Organisation named the disease Corona Virus Disease-2019 (COVID-19), and officially declared the pandemic state on March 11, 2020. Human coronaviruses have been known for decades as one of the causative agents of the common cold, but two previous coronavirus outbreaks, caused by the severe acute respiratory syndrome virus (SARS-CoV-1) and the Middle East respiratory syndrome virus (MERS-CoV), have demonstrated the remarkable pathogenic potential of human beta coronaviruses. Around 10,000 people have been infected by SARS and MERS, which has resulted in a death toll of about 1,500 patients, but the outbreaks remained largely confined in terms of time or spread, respectively. In contrast, SARS-CoV-2 spreads effectively and at a rapid pace by direct transmission with an R_0_ of at least 2 to 2.5 (2, 3). Due to high transmissibility and extensive community spread, this novel coronavirus has already caused over 12.1 million infections and over 550,000 deaths (as of 10 July 2020; https://www.who.int/emergencies/diseases/novel-coronavirus-2019), while world-wide shut-downs of social life and economy to confine the spread of this respiratory virus are causing have considerable impact.

After the emergence of SARS in 2002 and then MERS in 2012, vaccine development efforts have been initiated, including the use of recombinant measles virus (MeV) vaccine as a platform concept (4) to develop vector vaccine candidates against both agents and showed promising results. Recombinant MeV vectors encoding the unmodified SARS-CoV Spike protein induced high titers of neutralizing antibodies as well as IFN-γ T cell responses (5, 6) and conferred protection to immunized animals upon pathogen challenge by lowering virus titers more than 100-fold (5). For MERS, we have demonstrated that both, high titers of neutralizing antibodies as well as effective and polyfunctional T cell responses, were induced in vaccinated animals (7, 8) and conferred protection (7). Based on these data, a MeV-based MERS-vaccine candidate has been selected by the Coalition for Epidemic Preparedness Initiative (CEPI) for further clinical development (www.cepi.net/research_dev/our-portfolio).

Here, we explored the potential of recombinant MeV as vectors for the expression of the SARS-CoV-2 spike protein (S) as successfully applied for the development of MERS-(7, 8) and SARS-vaccine candidates (5, 6) as well as numerous other pathogens (4). The S glycoprotein was chosen as antigen for its role as primary target of neutralizing antibodies (6, 7) and the exemplary capability of MERS-CoV S protein to trigger strong cell-mediated immune responses when expressed by MeV in our front-runner MERS vaccine candidate (7, 8). The SARS-CoV-2 S protein-encoding gene was inserted into two different positions of the MeV genome to modulate antigen expression, and both recombinant MeV were successfully rescued. The virus expressing lower S protein levels resulted in stable amplification over at least 10 passages, while impairment of replication was insignificant. Indeed, immunization of IFNAR^-/-^-CD46Ge mice induced strong and functional humoral and cellular immune responses directed against both MeV and SARS-CoV-2 S protein biased for Th1-type T cell and antibody responses, illustrating the potential of MeV platform-based COVID-19 vaccine candidates.

## Results

### Generation and characterization of SARS-CoV-2-S by recombinant MeV_vac2_

Since the SARS-CoV and MERS-CoV spike proteins (S) have been shown to potently induce humoral and cellular immune responses, the SARS-CoV-2 S protein was chosen as appropriate antigen to be expressed by the recombinant MeV vaccine platform. A codon-optimized full-length gene encoding SARS-CoV-2 S protein was cloned into two different additional transcription units (ATUs) in the vaccine strain MeV_vac2_ genome, either downstream of the P (post P) or H (post H) gene cassettes (Fig. 1A). Recombinant viruses were successfully generated and amplified up to P10 in Vero cells with titers of up to 4×10^7^ TCID_50_/ml. The stability of the viral genomes was demonstrated via sequencing after RT-PCR of viruses in P2 or P10. In parallel to Sanger sequencing of the ATU-region encompassing the SARS-CoV-2-S gene, the full genome was sequenced using next generation sequencing with a coverage of 4 to 29,683 reads of each position (Suppl. Fig. S1) in P2. Both methods revealed no mutations across the whole vaccine genomes but a single A to G substitution on position 9 of the non-coding trailer region of the MeV_vac2_-SARS2-S(H) clone used for *in vivo* studies.

**Fig. 1:**
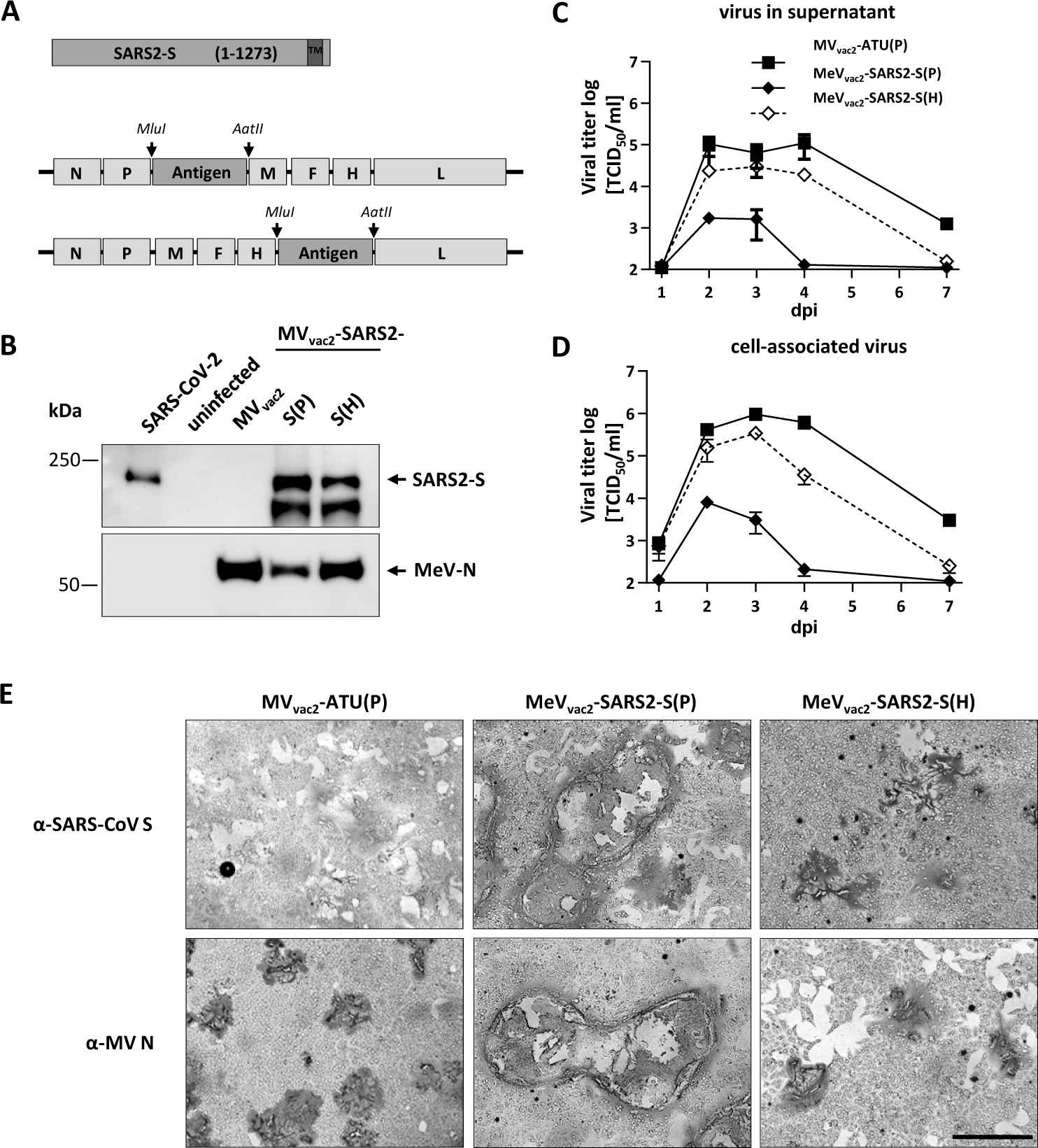
Generation and *in vitro* characterization of MeV_vac2_-SARS2-S(P) and MeV_vac2_-SARS2-S(H). **(A)** Schematic depiction of full-length SARS-CoV-2 S and recombinant MeV_vac2_ genomes used for expression of this antigen (lower schemes). Antigen or antigen encoding genes are depicted in dark grey; MeV viral gene cassettes (in light grey) are annotated. *MluI* and *AatII* restriction sites used for cloning of antigen-genes into post P or post H ATU are highlighted **(B)** Immunoblot analysis of Vero cells infected at an MOI of 0.01 with MeV_vac2_-SARS2-S(P), MeV_vac2_-SARS2-S(H), or MV_vac2_-ATU(P) (MV_vac2_) as depicted above lanes. Uninfected cells served as mock. Blots were probed using rabbit polyclonal anti-SARS spike antibody (upper blot) or mAb reactive against MeV-N (lower blot). Arrows indicate specific bands. (**C, D**) Growth kinetics of recombinant MeV on Vero cells infected at an MOI of 0.03 with MV_vac2_-ATU(P) or MeV_vac2_-SARS2-S encoding extra genes in post H or post P. Titers of samples prepared at indicated time points post infection were titrated on Vero cells. Means and standard deviations of three to five independent experiments are presented. (**E**) SARS-CoV-2 S protein expression in Vero cells was verified via immunoperoxidase monolayer assay. 50× magnification; scale bar, 500 μm.

To verify SARS-CoV-2 S protein expression levels, Western blot analysis of Vero cells infected with the MeV_vac2_-SARS2-S was performed. The S protein expression was slightly attenuated when cells were infected with viruses encoding the antigen in the ATU post-H compared to the post-P constructs (Fig. 1B). However, there was less overall viral protein expression in cells infected with post-P construct. Comparative growth kinetics with the vaccine viruses containing the SARS-CoV-2 S gene and the MV_vac2_-ATU(P) control virus revealed that the MeV_vac2_ encoding full-length SARS-CoV-2 S gene in post-P position grew remarkably different to the control virus, with approximately 100-fold reduced maximal titers. In contrast, growth of MeV_vac2_-SARS2-S(H) was much closer to MV_vac2_-ATU(P) with only a slight trend for lower titers (Fig. 1 C and D).

The impaired growth of MeV_vac2_-SARS2-S(P) was accompanied by a hyper-fusogenic phenotype (Fig. 1E, Suppl.Fig. S2A), which was also observed for the post-H vaccine candidate, but to a lesser extent. Therefore, fusion activity was quantified and compared to the parental MV_vac2_-ATU(P) as well as the MV_NSe_-GFP(N), which is known for its hyperfusogenic phenotype due to a V94M substitution in the F_2_ subunit of the MeV fusion protein (9). MV_vac2_-ATU(P) induced fusion of 16.8±0.8 (mean±SD) Vero cells 30 h after infection. MeV_vac2_-SARS2-S(P) revealed approximately 4-fold enhanced fusion activity (syncytia including 70±8 cells) while MeV_vac2_-SARS2-S(H) just fused 41±6 cells, thereby representing an intermediate phenotype. However, fusion activity of the latter became surpassed by MV_NSe_-GFP(N) that fused 56±4 cells in 30 h under the same conditions (Suppl. Fig. S2B).

To investigate if this increased fusion activity is due to SARS-CoV-2 S protein-mediated cell-to-cell fusion, we expressed the SARS-CoV-2 S protein by transfection of the eukaryotic expression plasmid pcDNA3.1-SARS2-S into SARS-CoV-2 receptor hACE2-negative 293T as well as into receptor-positive Vero cells. Indeed, expression of SARS-CoV-2 S protein induced syncytia of Vero, but not of 293T cells (Suppl. Fig. S3).

These data demonstrate that the hyperfusogenic phenotype of the SARS-CoV-2 S-encoding MeV is linked to expression of a fusion-active form of the SARS-CoV-2 S protein, indicating that cells infected by the vaccine candidates express a functional S protein. Thus, cloning and rescue of MeVs expressing correctly folded SARS-CoV-2 S was achieved successfully. Since higher S protein expression levels impaired viral replication, MeV_vac2_-SARS2-S(H) was chosen for further characterization *in vivo*.

### MeV_vac2_-SARS2-S(H) induces neutralizing antibodies against MeV and SARS-CoV-2

To test the efficacy of MeV_vac2_-SARS2-S(H) *in vivo*, genetically modified IFNAR^-/-^-CD46Ge mice were used, since they are the prime small animal model for analysis of MeV-derived vaccines (10). Groups of 4 -6 animals were immunized via the intraperitoneal (i.p.) route on days 0 and 28 with 1×10^5^ TCID_50_ of MeV_vac2_-SARS2-S(H) or empty MV_vac2_-ATU(P) as a control. As positive control, recombinant SARS-CoV-2 S protein adjuvanted with aluminum hydroxide gel (Alum) was injected subcutaneously, and medium-inoculated mice served as mock controls. 21 days after the second immunization, sera of immunized mice were analyzed in comparison to pre-bleed and post-prime immunization sera by ELISA on antigen-coated plates for total IgG antibodies binding to MeV bulk antigens (Fig. 2G-I) or SARS-CoV-2-S protein (Fig. 2A-C). Sera of mice vaccinated with MeV_vac2_-SARS2-S(H) contained IgG antibodies that bound to SARS-CoV-2-S protein (Fig. 2B and C), whereas no antibodies were found in mice before vaccination (Fig. 2A), or in MeV or mock-immunized control mice. Moreover, final sera of mice vaccinated with any recombinant MeV had IgG in the serum binding to MeV bulk antigens, indicating at least one successful vaccination with MeVs and general vector immunogenicity (Fig. 2G-I). The control S protein vaccine did induce higher levels of S protein-binding IgG than MeV_vac2_-SARS2-S(H).

**Fig. 2:**
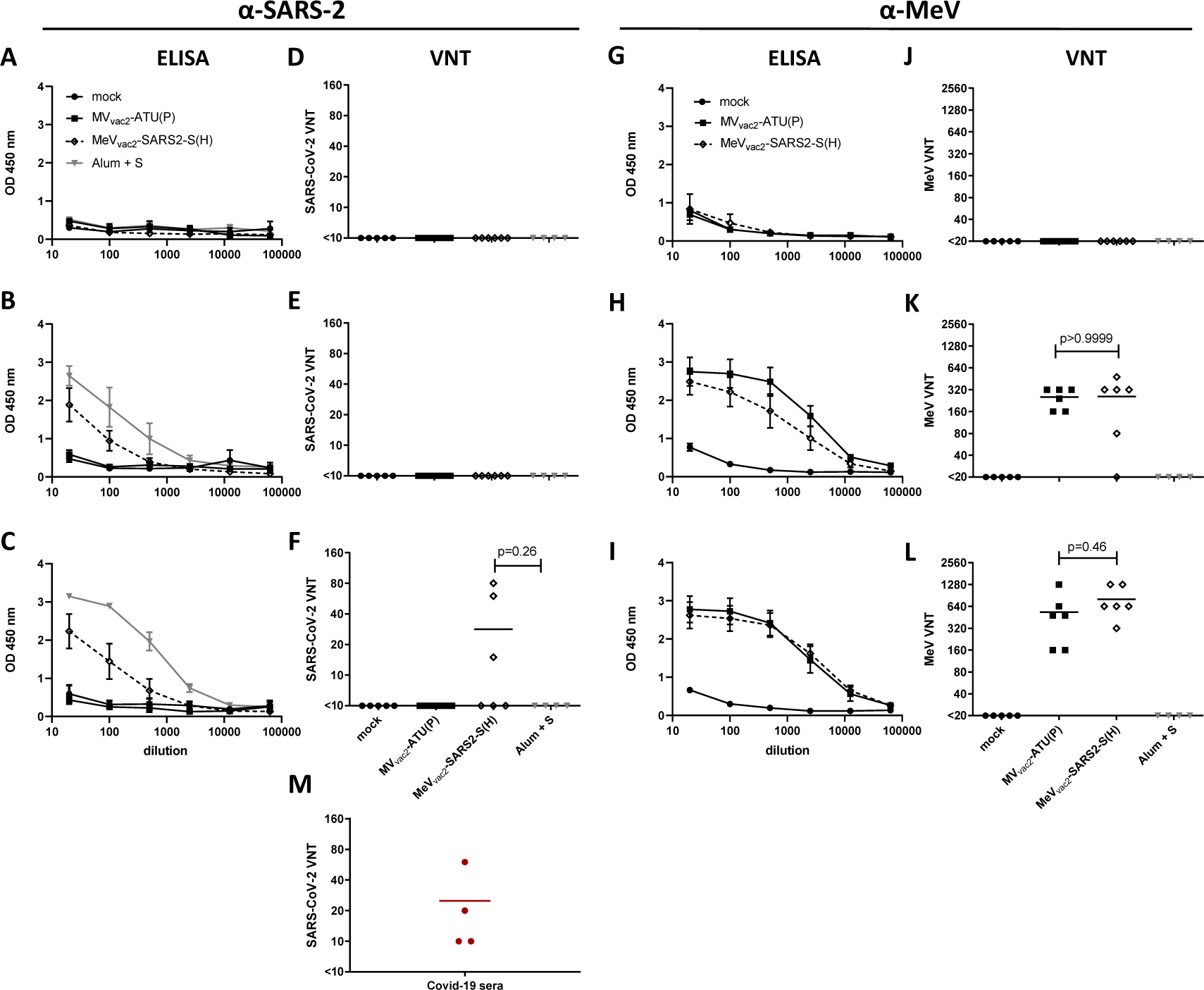
Induction of a-SARS-CoV-2 S and a-MeV specific antibodies. Sera of mice vaccinated on days 0 and 28 with indicated viruses or Alum-adjuvanted S protein were sampled on day 0 (A, D, E, F), day 28 after prime- (B, E, H, K) and day 49 after boost-immunization (C, F, I, L) and analyzed for antibodies specific for SARS-CoV-2 S or MeV. Medium-inoculated mice served as mock. Pan-IgG binding to recombinant SARS-CoV S (A – C) or MeV bulk antigens (G – I) were determined by ELISA via the specific OD 450 nm value. Depicted are means and respective standard deviation of the mean (SEM) of each group (n = 5 - 6). Virus neutralizing titers (VNT) in vaccinated mice for SARS-CoV-2 (D - F) or MeV (J – L) were calculated as reciprocal of the highest dilution abolishing infectivity. (**M)** SARS-CoV-2 VNT of 4 human Covid-19 reconvalescent sera. Dots represent single individuals; horizontal line represents mean per group. For statistical analysis of VNT data, one-way ANOVA was performed in combination with Tukey’s Multi comparison test to compare all pair means.

We next determined the neutralizing antibody responses against SARS-CoV-2 (Fig. 2D-F) or MeV (Fig. 2J-L). Most mice immunized with recombinant MeV, including those receiving the control virus, had developed MeV neutralizing antibody titers (VNT) after the first immunization (Fig. 2K). However, one mouse of the MeV_vac2_-SARS2-S(H) cohort initially reacted only weakly and another mouse not at all, reflecting individual differences in response to immunization. All animals had developed neutralizing antibodies after the second immunization, and a 3-fold increase was observed upon the second immunization (257 to 800 VNT, Fig. 2K, L). Neutralizing antibodies against SARS-CoV-2 were detected in mice vaccinated with MeV_vac2_-SARS2-S(H) after the second immunization, and reached a titer of 15 to 80 in three out of 6 mice (Fig. 2F). These titers were in the range of human reconvalescent sera tested in parallel (VNT 10 to 60; Fig. 2M). No VNTs against MeV or SARS-CoV-2 were detected in control mice inoculated with medium alone. Interestingly, the alumn-adjuvanted recombinant S protein did not induce any neutralizing antibodies despite higher binding IgG levels in ELISA, indicating that these antibodies bind to other epitopes of S or with lower affinity than those induced by the MeV-based vaccine candidate. In summary, the SARS-CoV-2-S protein-expressing MeV elicited robust neutralizing antibody responses against MeV and SARS-CoV-2.

### Splenocytes of animals vaccinated with MeV_vac2_-SARS2-S(H) react to SARS-CoV-2 S protein-specific stimulation

To assess the ability of MeV_vac2_-SARS2-S(H) to induce SARS-CoV-2-specific cellular immune responses, splenocytes of vaccinated animals were analyzed for antigen-specific IFN-γ secretion by ELISpot assay. Towards this, antigen-specific T cells were re-stimulated by co-cultivation with the syngeneic murine DC cell lines JAWSII or DC2.4 stably expressing the SARS-CoV-2 S protein. For JAWSII cells, bulk cultures of transduced cells were obtained by flow cytometric sorting. For DC2.4 cells, single cell clones were generated by limiting dilution of sorted respective of bulks cultures. Antigen expression by transduced DCs was verified by Western Blot analysis (data not shown).

ELISpot assays using splenocytes of vaccinated animals in co-culture with DC2.4-SARS2-S cells revealed more than 1,400 IFN-γ secreting cells per 1×10^6^ splenocytes after immunization with MeV_vac2_-SARS2-S(H), respectively (Fig. 3). In contrast, co-culture with splenocytes of control mice resulted in a background response of less than 50 IFN-γ producing cells per 1×10^6^ splenocytes. As expected, re-stimulation of T cells by DC2.4 presenting no exogenous antigen revealed only reactivity in the range of background (Fig. 3). To rule out clonal or cell line-associated artifacts, antigen-specific IFN-γ secretion by splenocytes of MeV_vac2_-SARS2-S(H) vaccinated mice was confirmed by stimulation with transgenic JAWSII-SARS2-S bulk cells. These cells also stimulated in excess of 1,400 IFN-γ secreting cells per 1×10^6^ splenocytes in animals receiving the recombinant SARS-CoV-2 vaccines, whereas only slight background stimulation was observed by the respective controls. The differences between MeV control and MeV_vac2_-SARS2-S vaccinated mice were statistically significant for both cell lines. Mice vaccinated with Alum-adjuvanted S protein showed no specific reactivity in IFN-γ ELISpot.

**Fig. 3:**
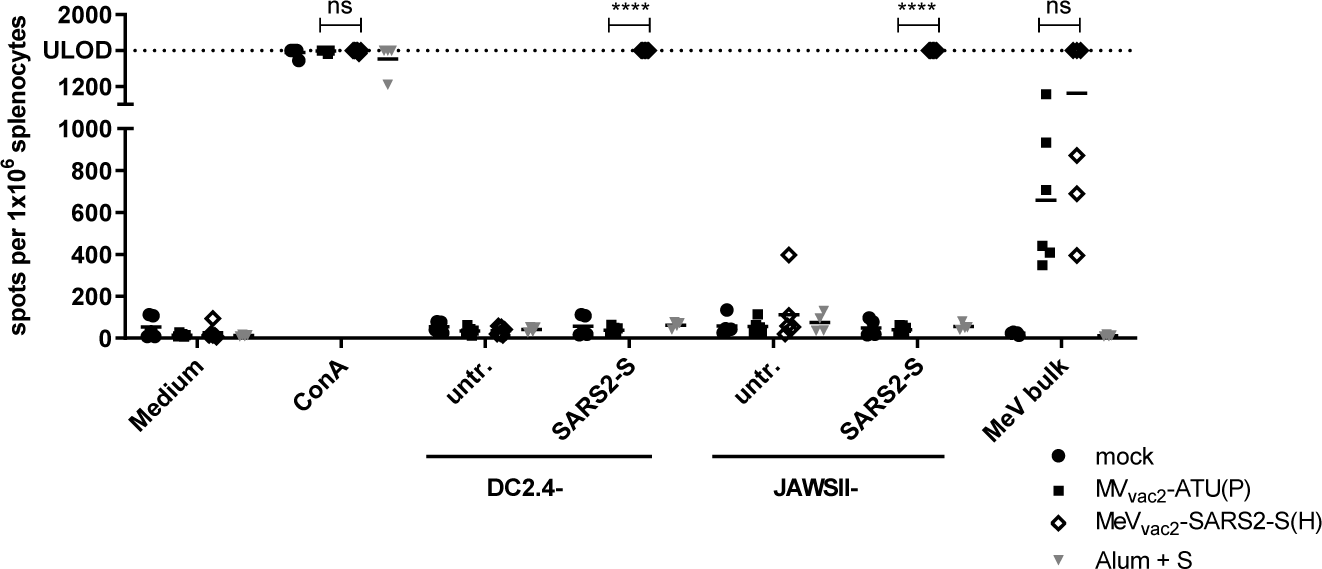
Secretion of IFN-γ after antigen-specific re-stimulation of splenocytes. IFN-γ ELISpot analysis using splenocytes of mice vaccinated on days 0 and 28 with indicated vaccines, isolated 21 days after boost immunization, and after co-culture with DC2.4 or JAWSII dendritic cell lines transgenic for SARS-CoV-2 S (SARS2-S) or untransduced controls (untr.). To analyze cellular responses directed against MeV, splenocytes were stimulated with 10 μg/mL MeV bulk antigens or were left unstimulated as controls (medium). The reactivity of splenocytes was confirmed by Concanavalin A (ConA) treatment (10 μg/mL). The number of cells per 1×10^6^ splenocytes represent the amount of cells expressing IFN-γ upon re-stimulation. Dots represent individual animals, horizontal bars mean per group (n = 5 - 6). Samples above the upper detection limit (ULOD) were displayed as such. For statistical analysis of grouped ELISpot data, two-way ANOVA analysis was applied with paired Tukey’s Multi comparison test used as post hoc test. ns, not significant (p>0.05); ****, p<0.0001.

Cellular immune responses upon stimulation with MeV bulk antigens were detected in animals that had been vaccinated with any recombinant MeV virus, as expected. While MeV bulk antigens stimulated only about 300 to 700 IFN-γ secreting cells per 1×10^6^ splenocytes of MV_vac2_-ATU(P) vaccinated animals, but 400 to 1,400 IFN-γ secreting cells per 1×10^6^ splenocytes of MeV_vac2_-SARS2-S(H) vaccinated animals. However, this trend was not statistically significant. Splenocytes of all animals revealed a similar basic reactivity to unspecific T cell stimulation, as confirmed by numbers of IFN-γ secreting cells upon ConA treatment at the limit of detection. Remarkably, stimulation of splenocytes by DC2.4 expressing SARS-CoV-2-S resulted in at least similar or even higher numbers of IFN-γ^+^ cells than after stimulation by MeV bulk antigens, indicating an extremely robust induction of cellular immunity against this antigen. Taken together, these data show that MeV_vac2_-SARS2-S(H) not only induces humoral, but also strong SARS-CoV-2-S protein-specific cellular immune responses.

### SARS-CoV-2 S-reactive T cells are multifunctional

To gain more detailed insights in the quality of the observed T cell responses, we further characterized the responsive T cell populations by flow cytometry, determining the expression of IFN-γ, TNF-α and IL-2 in CD8^+^ and CD4^+^ positive CD3^+^ T cells upon re-stimulation with SARS-CoV-2 S-presenting DC2.4-SARS2-S cells by intracellular cytokine staining (ICS). As a positive stimulus for T cell activation, tetradecanoylphorbol-acetate and ionomycin (TPA/Iono) were used. Exocytosis of cytokines was blocked by addition of brefeldin A (10 μg/mL) during stimulation. Cells were permeabilized, labelled, and fixed for flow cytometry. The gating strategy excluded duplicates (Suppl.Fig. S4, top row, middle panel), selected for living cells (Suppl. Fig. S4, top row, right panel), and separated CD8^+^ and CD4^+^ T cells on CD3^+^ cell populations (Suppl. Fig. S4, 2^nd^ row). Selected T cells were then analyzed for their expression of IFN-γ, TNF-α, or IL-2, double- (Suppl. Fig. S4, 3rd row), or triple-positive cells (Suppl. Fig. S4, bottom row) as exemplarily shown for CD4^+^ T cells after re-stimulation with TPA and ionomycin (Suppl. Fig. S4).

Vaccination with MeV_vac2_-SARS2-S(H) induced a significant amount of SARS-CoV-2 S-specific CD8^+^ T cells expressing either IFN-γ (Fig. 4B, left panel), IL-2 (Fig. 4B, middle panel) or TNF-α (Fig. 4B, right panel), with means between 0.1% and 0.5% of positive cells for each of these cytokines. Among those, a significant fraction of cells proved to be multifunctional, with a mean of 49% of the reactive CD8^+^ cells expressing two cytokines or 13% of responsive CD8+ cells being positive for TNF-α, IL-2 and IFN-γ (Fig. 4C). A much lower portion of responsive CD4^+^ T cells was observed, varying between 0.01% to 0.07% of CD4^+^ T cells. Among the responsive CD4+ cells, 46% expressed two cytokines and 10% were positive TNF-α, IL-2 and IFN-γ. Moreover, vaccination induced a significant fraction of vector-specific CD4^+^ T cells expressing IFN-γ (Fig. 4A, left panel), IL-2 (Fig. 4A, middle panel) or TNF-α (Fig. 4A, right panel) upon re-stimulation with MeV bulk antigen. Among those, multifunctional CD4^+^ T cells expressing two or all three cytokines were induced with a mean of about 22% and 6% poly-reactive T cells (Fig. 4C), respectively. To conclude, vaccination with MeV_vac2_-SARS2-S(H) induces not only IFN-γ, TNF-α, or IL-2 expressing T cells directed against SARS-CoV-2 and MeV, but also a significant fraction of multifunctional cytotoxic T cells specific for SARS-CoV-2 S and CD4^+^ T cells specific for MeV antigens, illustrating that a broad and robust SARS-CoV-2-specific immune response is induced by vaccination with MeV_vac2_-SARS2-S(H).

**Fig. 4:**
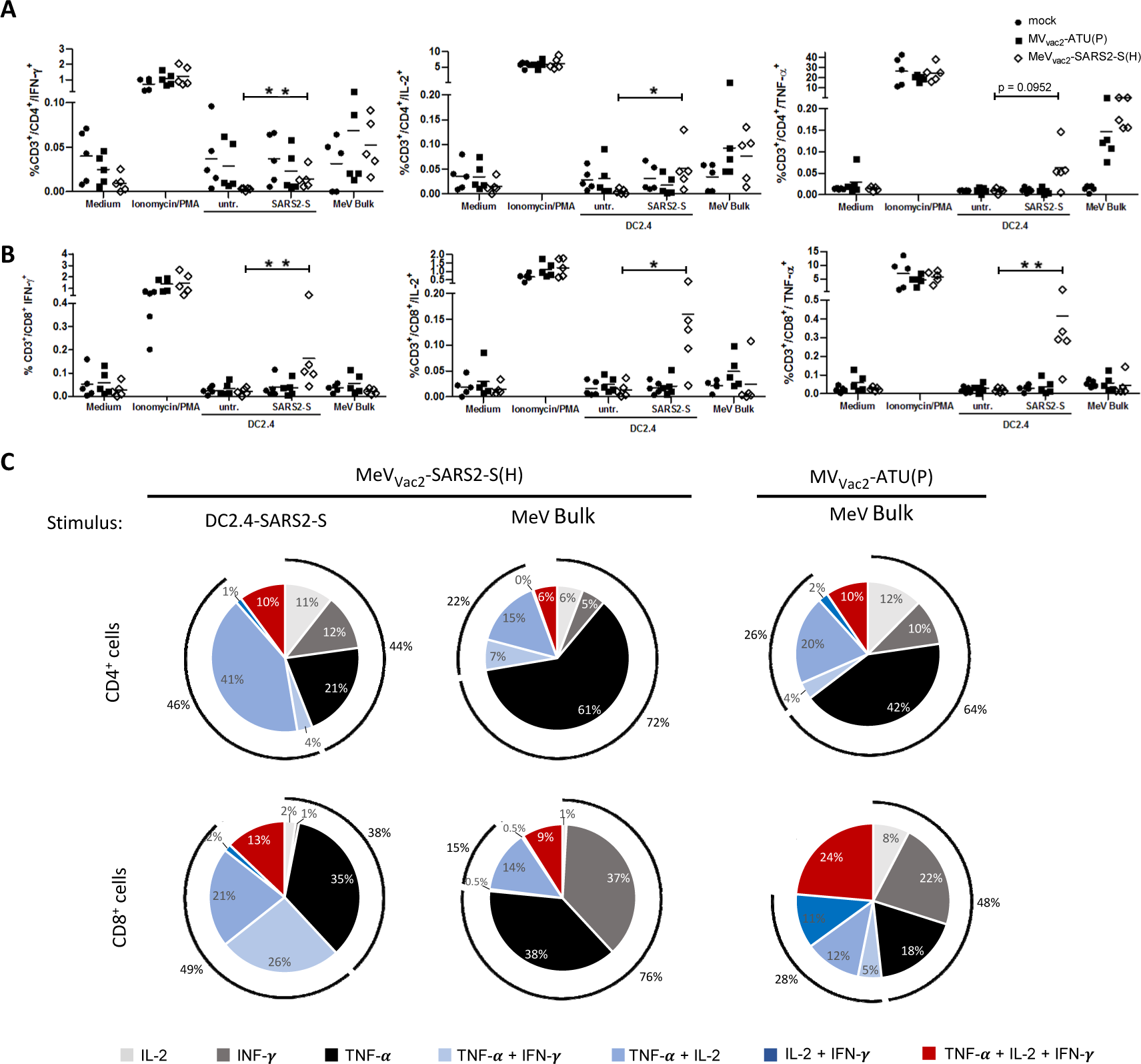
Detection of multi-functional T-cell responses induced by vaccination with MeV_vac2_-SARS2-S(H). Harvested splenocytes of MeV_vac2_-SARS2-S(H) vaccinated mice (same as depicted in Fig. 3) were re-stimulated and subjected to intracellular staining (ICS) for IFN-γ, TNF-α, and IL-2, and stained for extracellular T-cell markers CD3, CD4, and CD8 for flow cytometry analysis. Quantification of flow cytometry data of (**A**) CD4^+^ and (**D**) CD8^+^-positive T cells after co-culture with antigen-presenting DC2.4-SARS2-S or parental DC2.4 control cells, or after incubation with indicated stimuli (MeV bulk antigen (MeV bulk), or untreated cells (mock); reactivity of splenocytes was confirmed by ionomycin and phorbol myristate acetate (PMA) treatment (10 μg/mL). Dots represent individual animals, horizontal bars mean. Mann-Whitney test was used to compare cytokines levels between DC2.4 and DC2.4-SARS2-S re-stimulated splenoycytes in the MeV-_vac2_-SARS2-S(H) vaccine group without correction for multiple testing because of the exploratory character of the study. *, p<0.05; **, p<0.01. (**C**) reveals poly-functional T cells depicted in the pie-chart as fractions of cell populations expressing one, two, or all three of the tested cytokines and indicating the size of each fraction among all responsive T cells.

### MeV_vac2_-SARS2-S(H) induced antigen-specific CD8^+^ and CD4^+^ T cells respond with proliferation

While ELISpot and ICS analyses revealed antigen-specific cytokine secretion by vaccinated mice’ T cells, we next aimed at detecting antigen-specific CD8^+^ cytotoxic T lymphocytes (CTLs) which would be important for clearance of virus infected cell and CD4^+^ T helper cells. For that purpose, proliferation of CD8^+^ and CD4^+^ T cells upon stimulation with SARS-CoV-2-S was analyzed 3 weeks after the boost via a flow cytometry. Splenocytes of mice were isolated 21 days after the boost, and DC2.4-SARS2-S cells were used for re-stimulation of T cells. The splenocytes were labelled with CFSE and subsequently co-cultured with DC2.4-SARS2-S cells or, as a control, with parental DC2.4 cells for 6 days and finally stained for CD3, CD4, and CD8 before being analyzed for proliferation, detectable by the dilution of the CFSE stain due to cell division.

T cells of mice vaccinated with MeV_vac2_-SARS2-S(H) revealed an increase in the population of CD3^+^CD4^+^CFSE^low^ and CD3^+^CD8^+^CFSE^low^ cells after re-stimulation with DC2.4-SARS2-S cells compared to re-stimulation with parental DC2.4 without SARS-CoV-2 S antigen (Fig. 5). In contrast, T cells of control mice did not reveal this pattern, but the CFSE^low^ population remained rather constant. The prominent increase in CD3^+^CD8^+^CFSE^low^ cells, which was significant for MeV_vac2_-SARS2-S(H) vaccinated mice, indicates that CD3^+^CD8^+^ CTLs and CD3^+^CD4^+^ T helper lymphocytes specific for SARS-CoV-2 S have proliferated upon stimulation. Thus, SARS-CoV-2-specific cytotoxic memory T cells are induced in mice after vaccination with MeV_vac2_-SARS2-S(H).

**Fig. 5:**
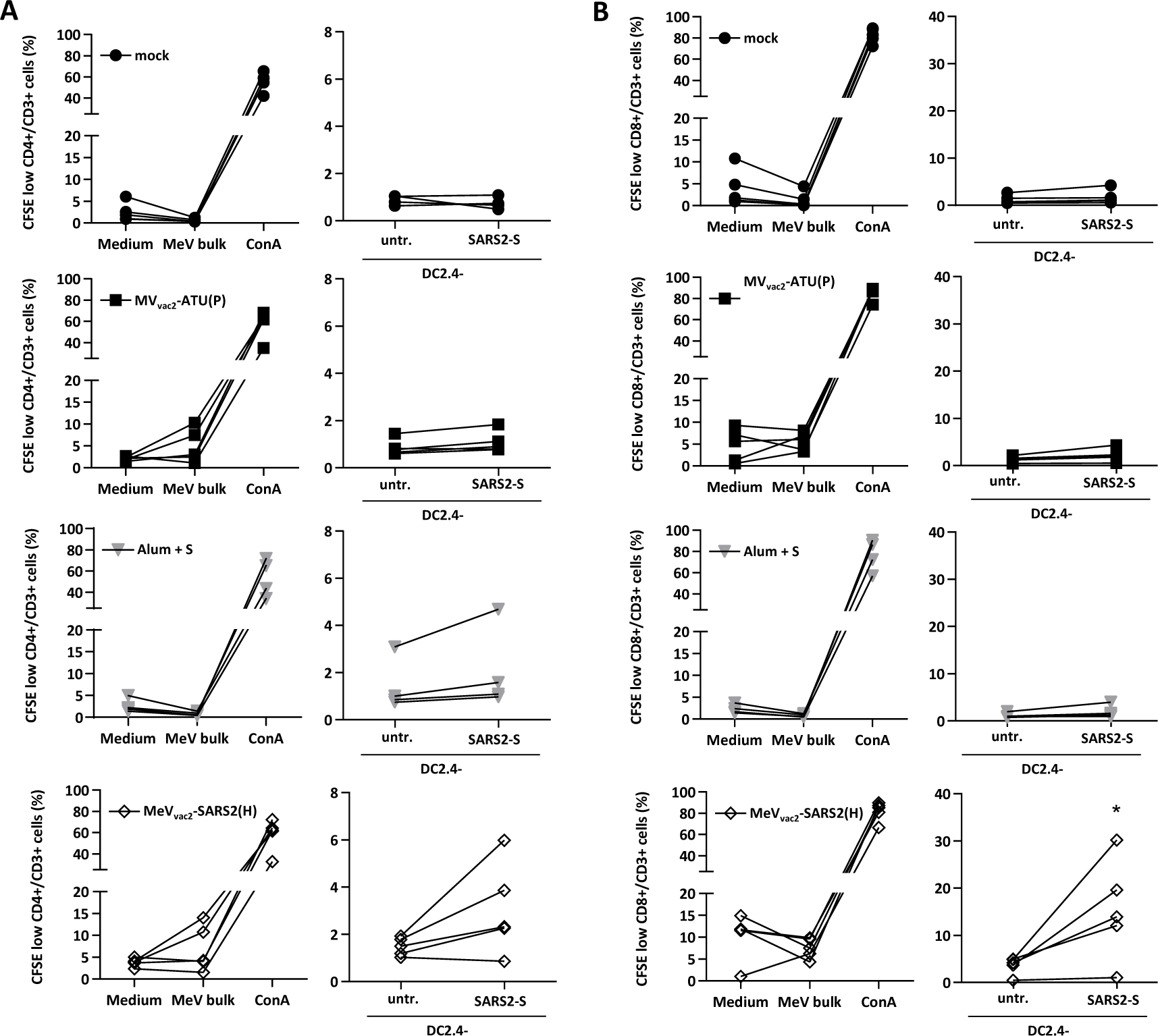
Ag-specific proliferation of SARS-CoV-2 S-specific T cells. Proliferation assay using splenocytes of mice vaccinated on days 0 and 28 with indicated viruses, isolated 21 days after boost immunization, after co-culture with DC2.4 dendritic cell line transgenic for SARS-CoV-2 S (SARS2-S) or untransduced parental DC2.4 (untr.). Depicted are the percentages of (**A**) CD4^+^ or (**B**) CD8^+^ T cells with low CFSE staining, indicating proliferation in the samples. To analyze cellular α-MeV responses, splenocytes were stimulated with 10 μg/ml MeV bulk antigens or were left unstimulated (medium). The reactivity of splenocytes was confirmed by concanavalin A (ConA) treatment (10 μg/ml). Results for splenocytes of vaccinated mice are displayed individually and the trend between paired unstimulated and re-stimulated samples is outlined (n = 2-4). One-tailed Mann-Whitney t-test. *, p<0.05.

### Induced T cells reveal antigen-specific cytotoxicity

To demonstrate the effector ability of induced cytotoxic T lymphocytes (CTLs), a killing assay was performed to directly analyze antigen-specific cytotoxicity (Fig. 6). Splenocytes of immunized mice isolated 21 days post boost vaccination were co-cultured with DC2.4-SARS2-S or parental DC2.4 cells for 6 days to re-stimulate antigen-specific T cells. When these re-stimulated T cells were co-incubated with a defined mixture of EL-4_green_-SARS2-S target and EL-4_red_ control cells (ratio approximately 1:1), only T cells from MeV_vac2_-SARS2-S(H) vaccinated mice significantly shifted the ratio of live SARS-CoV-2 S protein-expressing target cells to control cells in a dose-dependent manner (Fig. 6B). This antigen-dependent killing was also dependent on re-stimulation with DC2.4-SARS2-S cells, since unstimulated T cells did not significantly shift the ratios of target to non-target cells (Fig. 6A).

**Fig. 6:**
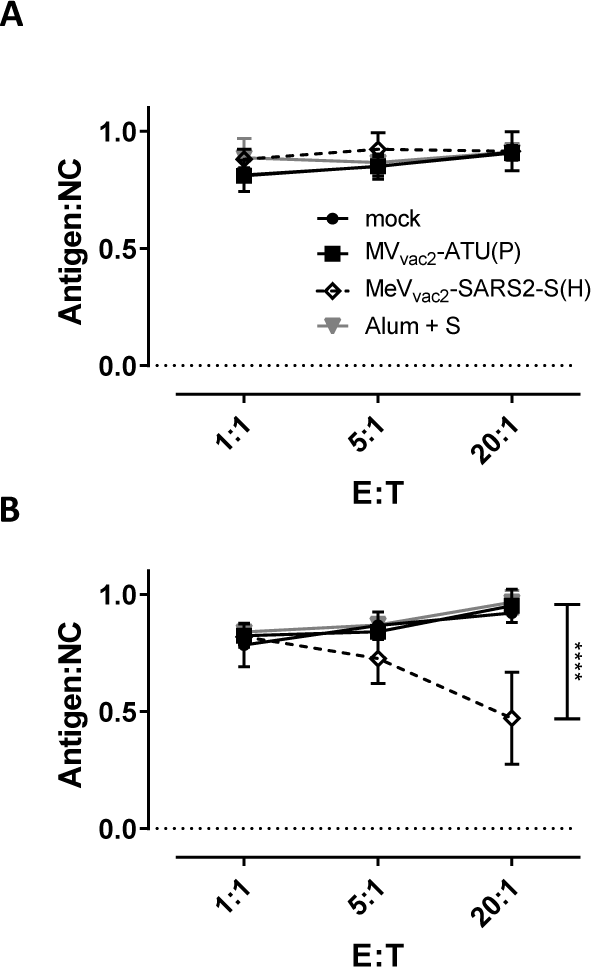
Antigen-specific killing activity of SARS-CoV-2 S-specific T cells. Killing assay using splenocytes of mice vaccinated on days 0 and 28 isolated 21 days after the second immunization. Splenocytes were co-cultured with DC2.4 (**A**) or with antigen-presenting DC2.4-SARS2-S (**B**) cells or for 6 days. Activated CTLs were then co-cultured with EL-4_green_-SARS2-S target cells (Antigen) and EL-4_red_ control cells (NC) at indicated E:T ratios for 4 h. Ratio of living target to non-target cells (Antigen:NC) was determined by flow cytometry. Depicted are means and standard deviation of each group (open diamonds, MeV_vac2_-SARS2-S(H); filled circles, mock; filled squares, MV_vac2_-ATU(P); grey triangles: S protein + Alum) (n = 3 - 5). For statistical analysis of grouped ELISpot data, two-way ANOVA analysis was applied with paired Tukey’s Multi comparison test used as post hoc test. ****, p<0.0001.

These results indicate that CTLs isolated from MeV_vac2_-SARS2-S(H)-vaccinated mice are capable of lysing cells expressing SARS-CoV-2 S. Neither splenocytes of control mice re-stimulated with DC2.4-SARS2-S nor splenocytes of SARS-CoV-2-S vaccinated mice re-stimulated with control DC2.4 cells showed such an antigen-specific killing activity, demonstrating that MeV_vac2_-SARS2-S(H) induces fully functional antigen-specific CD8^+^ CTLs.

### Induced immunity is skewed towards Th1-biased responses

While the functionality of both humoral and cellular anti-SARS-CoV-2 immune responses elicited by MeV_vac2_-SARS2-S(H) is reassuring, the SARS-CoV-2 vaccine development has to proceed with some caution because of the potential risk of immunopathogenesis observed in some animal models, such as antibody-dependent enhancement (ADE) and enhanced respiratory disease (ERD) which seem to correlate with a Th2-biased immune response. Since in mice IgG1 is a marker for Th2-bias and risk of ADE development, whereas IgG2a antibodies indicate a favorable Th1-bias, IgG subtype-specific ELISA were performed with the sera collected at different time points. Animals vaccinated with alum-adjuvanted SARS-CoV-2 S protein, a vaccine concept known for its Th2-bias (11, 12), developed high levels of S protein-specific IgG1 antibodies, whereas few S-specific IgG2a antibodies were detected (Fig. 7A). In comparison, MeV_vac2_-SARS2-S(H) induced 100-fold less IgG1 antibodies, but at least 10-fold higher IgG2a levels (Fig. 7A), indicating a favorable Th1-bias in animals immunized with the MeV-derived vaccine candidate.

**Figure 7:**
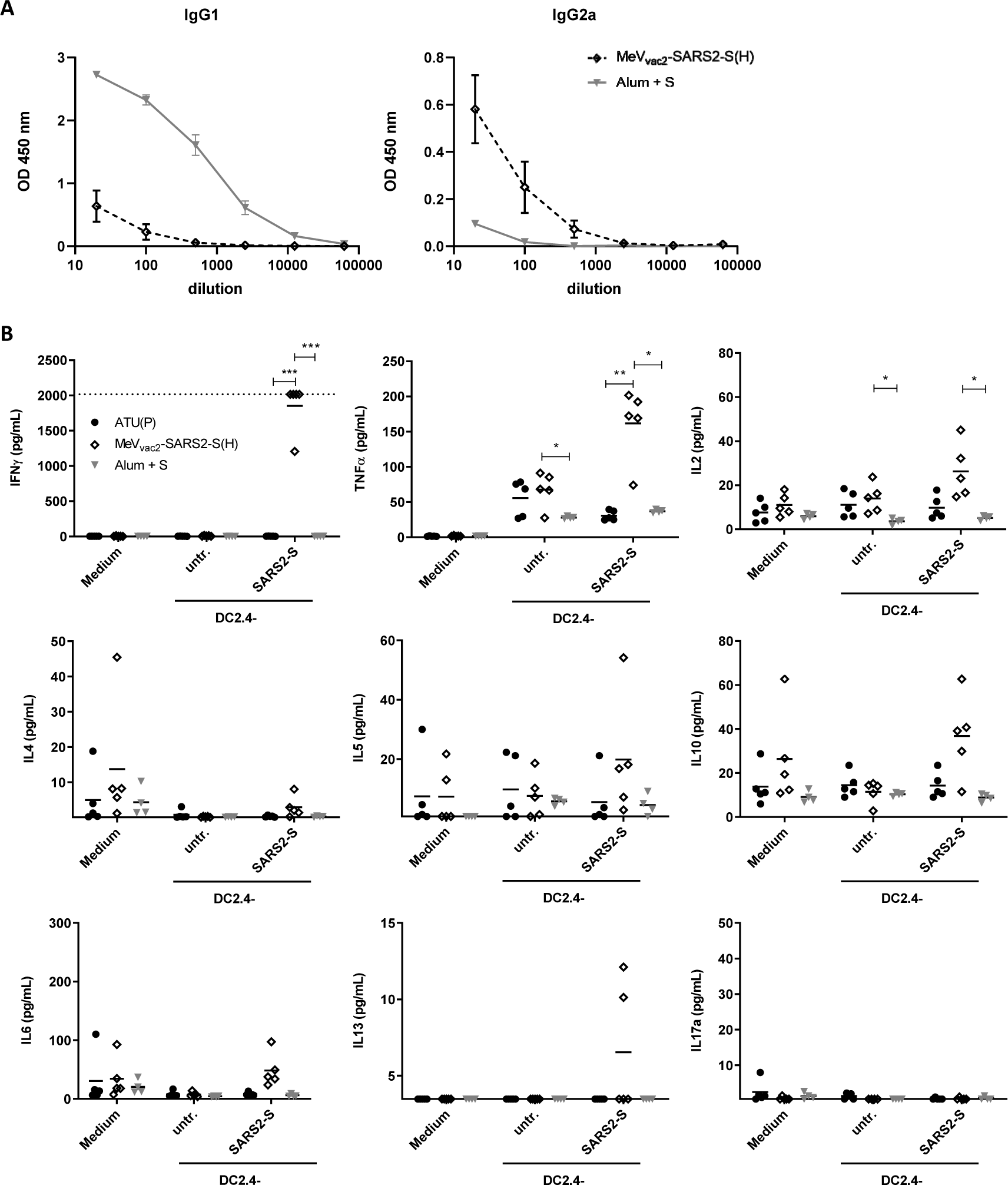
Immune bias of induced responses. To analyze skewing of immune responses towards Th1- or Th2-biased immunity (**A**) sera and (**B**) splenocytes of vaccinated mice depicted before were analyzed. (**A**) Sera of mice vaccinated on days 0 and 28 with MeV_vac2_-SARS2-S(H) or Alum-adjuvanted S protein already shown in Fig. 2 were analysed for IgG1- or IgG2a-type antibodies specific for SARS-CoV-2 S. IgG1 (left panel) or IgG2a (right panel) binding to recombinant SARS-CoV S were determined by ELISA via the specific OD 450 nm value. Depicted are means and respective standard deviation of the mean (SEM) of each group (n = 5 - 6). (**B**) Splenocytes of the same mice already shown in Figs. 3 to 6 were analysed by multiplex cytokine analysis for secretion of typical marker cytokines in the supernatant after re-stimulation by co-culture with antigen-presenting DC2.4-SARS2-S cells. DC2.4 cells served as non-specific control stimulus. Dots represent individual animals, horizontal bars mean per group (n = 4 - 5). IFN-γ: upper limit of detection (ULOD): 2015.2 pg/mL; IL-6: ULOD: 3992,4 pg/mL; IL-17a lower limit of detection (LLOD): 0.473 pg/mL; IL-4 LLOD: 0.095 pg/mL; IL-5 LLOD: 0.685 pg/mL; IL-13 LLOD: 3.463 pg/mL. For statistical analysis of grouped multiplex data, two-way ANOVA analysis was applied with paired Tukey’s Multi comparison test as post hoc test. *, p<0.05; **, p<0.01; ***, p<0.00

These findings were confirmed by multiplex cytokine analysis of the cytokine profile in the supernatants of splenocytes from vaccinated animals, which were re-stimulated using DC2.4 or DC2.4-SARS2-S cells. All splenocytes revealed secretion of all cytokines after stimulation with ConA demonstrating general reactivity of cells and assay (data not shown). Most likely due to the low number of S-reactive T cells in animals that had been vaccinated with recombinant SARS2-S protein and Alum, no or minimal, constant cytokine levels were measurable in the supernatants of re-stimulated splenocytes (Fig. 7B). In contrast, splenocytes of animals immunized with MeV_vac2_-SARS2-S(H) reacted specifically with the secretion of IFN-γ, TNF-α, and IL-2 upon re-stimulation by DC2.4-SARS2-S (Fig. 7B, top row), in accordance with ELISpot and ICS data. However, we could observe no or minimal up-regulation of IL-4, IL-5, IL-13, or IL-10, which would have been indicative for a Th2-biased response (Fig. 7B, middle row). Also IL-17a, or IL-6 indicative of a Th17 or general inflammatory response showed minimal changes (Fig. 7 B, bottom row).

Thus, both humoral and cellular responses reveal a Th1-biased immunity induced by MeV_vac2_-SARS2-S(H), which indicates a relatively low risk for putatively Th2-mediated immunopathologies.

## Discussion

In this study, we aimed to analyze the efficacy of MeV-derived vaccine candidates encoding the Spike glycoprotein S of SARS-CoV-2 to induce functional immune responses to protect against COVID-19. We show that MeV_vac2_-SARS2-S(H) replicated comparably to MeV vaccine strain viruses and was genetically stable over extended passaging. Upon vaccination of mice, it induced robust humoral immune responses of the IgG2a subtype directed against the SARS-CoV-2 spike glycoprotein S with neutralizing activity in a range already shown to be protective by others. In addition, considerable amounts of SARS-CoV-2 S-specific CD4^+^ and CD8^+^ T cells were induced, the major fraction of which were secreting two or even all three cytokines when analysing for IFN-γ, TNF-α, or IL-2 upon antigen-specific re-stimulation. These T cells proliferated and specifically depleted antigen-positive target cells in a mixed population. Importantly, all responses were skewed toward Th1-biased immunity. In parallel, the capacity to induce measles-specific immune reactivity remained conserved.

This effective MeV Moraten strain-derived recombinant vaccine MeV_vac2_-SARS2-S(H) is a live-attenuated vaccine that encodes the full-length, functional version of the SARS-CoV-2 Spike protein as main target for functional antibodies, but also for induction of T cell responses. Vero cells revealed homogenous expression of the SARS-CoV-2 S antigen by Western Blot analyses and positive immunostaining of syncytia after infection by MeV_vac2_-SARS2-S(H). Stable antigen expression is a prerequisite for the immune system to encounter the specific antigen to mount robust immune responses and for industrial production of a vaccine. Indeed, IFNAR^-/-^-CD46Ge mice vaccinated with MeV_vac2_-SARS2-S(H) in a prime-boost protocol showed uniform induction of antibodies directed against MeV bulk antigens or SARS-CoV-2 S, which had considerable neutralizing activity against both pathogens. We observe antibody responses in these animals at a level that correlate with protection in mouse challenge models (13), as well as with neutralizing activity we found in the serum of 4 reconvalescent human patients. These responses were triggered even though the knock-out of the type I interferon receptor, which is necessary to allow propagation of MeV in mice (10, 14). This knock-out usually should impair the induction of especially humoral immune responses (15). This highlights the remarkable immunogenicity of the MeV vaccine platform technology that also works in this model with partially impaired immune responses.

However, why did not all immunized animals develop neutralizing activity detectable in our assay? Firstly, determination of the VNT relying on 100% pathogen neutralization is obviously a rather harsh assay in the context of SARS-CoV-2, as evidenced by the modest VNT titers published so far, in general, and absence of VNT in the S+Alum vaccinated group despite high amounts of S binding antibodies. This means that just detectable VNT already indicates considerable neutralizing activity. Secondly, we realized that two of the three animals which did not show a VNT >10 have not responded well to the prime vaccination, at all. These animals developed none or only a minor VNT against MeV after the first vaccination. This observation is rather unusual, and argues for technical issues during the first vaccination in these two animals. Since none of the animals showed VNT against SARS-CoV-2 after one vaccination with the vaccine, it is tempting to speculate that a prime-boost-protocol is associated in this animal model with maturation of antibodies to generate better neutralizing responses. On the other hand, all animals including the two improperly immunized ones revealed significant, multi-functional T cell responses against SARS-CoV-2 S, which were still recordable three weeks after the second vaccination, when we already expect constriction of antigen-specific T cell effector populations. These data suggest that anti-S antibody responses mature after repeated vaccination, but on the other hand that a one shot vaccination regime will already induce especially functional memory T cell immune responses, the protective efficacy of which as well as their duration has to be demonstrated in future challenge experiments. In any case, we have observed with other foreign viral antigens that these T cell responses can be detected in mice more than 2 years after vaccination (Hörner & Fiedler et al., unpublished data). This observation is in accordance with the stability of anti-measles immunity (16) also after pediatric vaccination (17) and might be a specific advantage of the measles vaccine platform technology. Also extended passaging of the vaccine candidate did not result in changes of the vaccine as revealed by sequencing of the virus after 10 passages starting with a low MOI. This genetic stability indicates that the slight impairment seen in multi-step growth curves when compared to a vaccine-strain MeV is not critical for the vaccine’s amplification and therefore crucial for product safety. In accordance with its genetic stability, the minor enhancement of fusion activity can also be regarded as non-critical, especially with a view on the fusion activity of MeV used in clinical trials for treatment of tumors. These so called oncolytic MeV have been used in 15 phase 1 and phase 2 clinical trials, so far. Thereby, advanced-stage tumor patients suffering from different tumor entities have been treated. Despite constituting in principle a vulnerable patient collective, application of high doses of non-targeted, fusion-active MeV (up to 1×10^11^ TCID_50_) (18) systemically or for example directly into the patients’ brains (19) was accompanied by an acceptable safety profile (20). Therefore, the enhancement of fusion activity cannot be expected to be crucial for product safety, while the attenuation of vaccine-strain MeV is multifactorial, anyway, and not just a matter of cell entry tropism and mechanism (21). Likewise, the clinical phase 1 and 2 trials using the MeV vector platform for the generation of bivalent vaccines, which induce immunity against CHIKV (22, 23), have revealed an extremely beneficial safety profile of this recombinant vaccine concept also in human patients, while signs of efficacy became evident.

In any case, generation of MeV-derived COVID-19 vaccines encoding a less fusion-active variant of the SARS-CoV-2 Spike glycoprotein might be beneficial to enhance titers of the vaccine virus. In the meantime, stabilized S variants have become available that have attenuated or no cell-cell fusion activity. One variant has a deletion of the multi-basic cleavage motif for furin-like proteases at the S1/S2 boundary that facilitates pre-activation of S (24). A second variant has proline substitutions at residues 986 and 987, which are stabilizing a pre-fusion conformation of S (25). Vaccine candidates encoding S with one of these motifs or a combination thereof in a soluble version as already done for DNA vaccines (26) are under development. These have to show an at least comparable capacity to induce neutralizing antibody responses also in the context of MeV infection, which might be dependent on the respective conformation of the antigen that is expressed by vaccine virus-infected cells *in situ*.

The induction of the “right” antibodies and T cell responses is especially crucial with a view on potential complications that can be observed when coronavirus encounter “wrong” immune responses that can give rise to immunopathologies after infection. In some infected cats, infection with feline coronavirus causes feline infectious peritonitis, a deadly disease characterized by viral infection of macrophages during the acute phase. Interestingly, the switch of pathology after infection from a rather moderate pathogenesis into an acute, devastating disease can be triggered by vaccination of persistently infected cats and has been attributed to the induction of antibodies that mediate enhancement of the disease, a process called antibody-dependent enhancement (ADE). During COVID-19, ADE might be the cause of the severe cases currently observed. Some case reports indicate that severe disease appeared more frequently in patients with high SARS-CoV-2 immunoglobulin G (IgG) levels (27). ADE has been most prominent for dengue virus (DENV) infections, especially in secondary infections with a different DENV serotype where enhancement of disease correlated with the induction of non-neutralizing Abs that can mediate an efficient uptake of the virus in FcR-positive cells such as macrophages and other immune cells (28). Moreover, other immune-related adverse events were described for SARS- and MERS-CoV. When animals were immunized with vaccines that pre-dominantly induce Th2-biased T-helper cell responses, vaccinated mice revealed significantly reduced virus loads after challenge, but also an eosinophilic infiltrate into the lungs accompanied by pathological changes of the lung tissue, so called enhanced respiratory disease (ERD) (29). Such immunopathologies upon CoV infection are a major concern for diseases pathology and especially vaccine development. Thus, Th2-biased immune responses as triggered by alum-adjuvanted whole inactivated virus particles or recombinant proteins should be avoided.

Interestingly, the live-attenuated MeV vaccine is known for a balanced Th1/Th2-bias of induced immune responses with a bias for Th1 responses at least during the acute phase after vaccination (30). In theory, this should also apply for immune responses induced against all antigens presented during a MeV vaccine virus infection including foreign antigen(s) additionally expressed when MeV is used as vaccine platform. Indeed, our analyses provide evidence that the bias of the immune responses is in favour of Th1 responses, as revealed by the inverted IgG1/IgG2a subtype ratio of antibodies induced against SARS-CoV-2 S by MeV_vac2_-SARS2-S(H) compared to the animals immunized with alum-adjuvanted recombinant S protein. Moreover, the cytokine profile of splenocyte cultures of immunized mice after re-stimulation of S-specific T cells reveals a respective preferable Th1 bias. Since SARS-CoV-2 and SARS-CoV use the same primary attachment receptor for cell entry, hACE2, and selected hACE2-transgenic mice show differential pathology after inoculation with SARS-CoV-2 (13, 31), studying the impact of the Th1-biased MeV-based immunization in hACE2-transgenic mice during challenge with SARS-CoV-2 will be a matter of future studies. In any case, our data suggest that MeV-derived COVID-19 vaccines have a low likelihood to trigger immunopathogenesis. Another animal model for COVID-19, golden Syrian hamsters, could be an alternative for future challenge studies. This animal model is susceptible for SARS-CoV-2 infection, reveals a moderate, but clearly distinguishable pathology, and shows air-borne transmissibility from infected to naïve animals (32, 33). Therefore, this animal model accurately reflects at least some aspects of the course of human disease and should be valuable for assessment of the protective efficacy of COVID-19 vaccines.

In conclusion, the bivalent MeV/SARS-CoV-2 vaccine candidate has a number of desirable properties with respect to its immunogenicity against SARS-CoV-2. Furthermore, the concurrent induction of anti-MeV immunity would allow its use in the context of routine measles immunization schedules. Such a MeV-based COVID-19 vaccine could be included in the currently applied MMR (measles, mumps, rubella) vaccine, providing additional protection against SARS-CoV-2. While controversially discussed to which extent, children do become infected and shed the virus, despite them rarely being severely affected. In any case, preventing infection or virus shedding from vaccinated children can also help to contain the disease and protect vulnerable patient groups. Moreover, the capacity to produce large amounts of vaccine doses would be available more or less instantly from routine measles vaccine production, but at no impairment of production of other necessary vaccines, since the measles vaccine property is preserved in the proposed vaccine candidate. Especially since vaccination against the measles should not be impaired also during the COVID-19 epidemic, this is a considerable advantage. Otherwise, parallel epidemics with another, even more contagious respiratory virus are looming when vaccination programs against the measles are stopped in favour of COVID-19 vaccination programs. Therefore, MeV_vac2_-SARS2-S(H) is a promising vaccine candidate that warrants further investigation.

## Materials and Methods

### Cells

Vero (African green monkey kidney) (ATCC# CCL-81), Vero clone E6 (ATCC# CRL-1586), 293T (ATCC CRL-3216) and EL-4 (ATCC TIB-39) cell lines were purchased from ATCC (Manassas, VA, USA) and cultured in Dulbecco’s modified Eagle’s medium (DMEM, Biowest, Nuaillé, France) supplemented with 10% fetal bovine serum (FBS; Biochrom, Berlin, Germany) and 2 mM L-glutamine (L-Gln; Biochrom). JAWSII mouse dendritic cells (ATCC CRL-11904) were also purchased from ATCC and cultured in MEM-α (GIBCO BRL, Eggenstein, Germany) supplemented with 20% FBS, 2 mM L-Gln, 1 mM sodium pyruvate (Biochrom), and 5 ng/ml murine GM-CSF (Biotechne, Wiesbaden, Germany). DC2.4 mouse dendritic cells (34) were cultured in RPMI containing 10% FBS, 2 mM L-Gln, 1% non-essential amino acids (Biochrom), 10 mM HEPES (pH 7,4), and 50 μM 2-mercaptoethanol (Sigma-Aldrich, Steinheim, Germany). All cells were cultured at 37°C in a humidified atmosphere containing 6% CO_2_ for a maximum of 6 months of culture after thawing of the original stock.

### Plasmids

The codon-optimized gene encoding full-length SARS-CoV-2 Spike glycoprotein S of isolate Wuhan-Hu-1 (Genebank accession no. MN908947.1) in plasmids pMA-RQ-SARS2-S flanked with *Aat*II/*Mlu*I and *NheI*/*XhoI* restriction sites was obtained by gene synthesis (Invitrogen Life Technology, Regensburg, Germany). The antigen was inserted into plasmids pBRPolIIΔ-MV_vac2_-GFP(P) or pBRPolIIΔMV_vac2_-GFP(H) via *Mlu*I/*Aat*II to generate pBRPolII-MV_vac2_-SARS2-S(P) or pBRPolII-MV_vac2_-SARS2-S(H). pBRPolIIΔ-MV_vac2_-GFP(P) or pBRPolIIΔMV_vac2_-GFP(H) were generated by inserting the immediate early CMV promoter sequence from p(+)PolII-MV_NSe_-GFP(N) (35), which had been modified by site-directed mutagenesis for deleting the *AatII* restriction sites, into pBR-MV_vac2_-GFP(P) or pBR-MV_vac2_-GFP(H) (7). For construction of a lentiviral transfer vector encoding SARS-CoV-2 S directly linked to the *egfp* gene as selection marker, the ORF of SARS-CoV-2 S was inserted via *Nhe*I/*Xho*I into pCSCW2gluc-IRES-GFP (36) to yield pCSCW2-SARS2-S-IRES-GFP. For construction of a eukaryotic expression plasmid encoding SARS-CoV-2-S, the ORF of SARS2-S was inserted via *Nhe*I/*Xho*I into pcDNA3.1(+) (Invitrogen Life Technology) to yield pcDNA3.1-SARS2-S.

### Production of lentiviral vectors and generation of antigen-expressing dendritic cell lines

Lentiviral vectors were produced and used for the generation of antigen-expressing dendritic cell lines as described before (7). In short, HIV-1-derived particles pseudotyped with VSV-G were generated using a standard three plasmid system, pMD2.G, pCMVΔR8.9 (37) with the transfer vector plasmid pCSCW2-SARS2-S-IRES-GFP in combination with PEI transfection of 293T cells (38). Subsequent purification by filtration and ultracentrifugation of supernatants yielded virus stocks were used to transduce murine DC cell lines, DC2.4 and JAWSII, as well as the murine T cell line EL-4, resulting in DC2.4-SARS2-S, JAWSII-SARS2-S, and EL-4_green_-SARS2-S, respectively, that express the SARS-CoV-2 S protein and GFP and present the respective peptides via MHC-I. Transduced cultures with 1-10% GFP-positive cells were single cell-sorted (BD FACS Aria™ Fusion) for GFP-expressing cells and subsequently characterized for antigen expression. For JAWSII-SARS2-S, the bulk-sorted cells were used in stimulation experiments. For DC2.4-SARS2-S and EL-4_green_-SARS2-S, clonal cell lines were generated by limiting dilution of bulk-sorted cells and characterized for marker- and antigen-expression.

### Viruses

SARS-CoV-2 S-encoding vaccine candidates MeV_vac2_-SARS2-S(P) or MeV_vac2_-SARS2-S(H) were generated as described previously (7, 39). Single syncytia were picked and overlaid onto 50% confluent Vero cells cultured in 6-well plates and harvested as “passage 0” (P0) by scraping and freeze-thaw cycle of cells at the time of maximal infection. Subsequent passages were generated after TCID_50_ titration of infectious virus according to the method of Kaerber and Spaerman (40). Stocks were generated by infection of Vero cells at an MOI = 0.03, and passage 2 (P2) or P3 were used for *in vitro* characterization, while vaccine viruses in P3 or P4 were used for vaccination experiments. Vector control virus MV_vac2_-ATU(P) (41) was used in P5 for vaccination. SARS-CoV-2 (isolate MUC-IMB1) (kind gift of G. Dobler, Bundeswehr Insitute for Microbiology, Germany) was used for SARS-CoV-2 neutralization assays. It was propagated on Vero E6 cells and was titrated via TCID_50_ as described above for recombinant MeV. All virus stocks were stored in aliquots at - 80°C.

Multistep viral growth kinetics were analyed by infecting Vero cells at an MOI of 0.03 in 96-well plates and incubated at 37°C. At various time points, supernatants were clarified by centrifugation, and cells were scraped into OptiMEM and subjected to freeze-thaw cycles. Released and cell-associated viral titers were determined by TCID_50_ limited dilution method.

### Measles virus genome sequence analysis

The RNA genomes of recombinant MeV in P2 or P10 were isolated from infected Vero cells using the QIAamp Viral RNA Mini Kit (QIAgen, Hilden, Germany) according to the manufacturer’s instructions and resuspended in 50 μL RNase-free water. Viral cDNA was reversely transcribed using Superscript II RT kit (Invitrogen) with 2 μL viral RNA as template and random hexamer primers, according to manufacturer’s instructions. For specific amplification of the SARS-CoV-2 S ORF, the respective genomic regions of recombinant MeV were amplified by PCR using primers binding to sequences flanking the regions of interest and the cDNA as template. Detailed description of primers and procedures are available upon request. The PCR products were directly sequenced (Eurofins Genomics, Ebersberg, Germany).

### NGS library preparation and sequencing

Total RNA was isolated from Vero cells after 4 days post infection using the Direct-zol RNA isolation kit (Zymo Research). 1 μg of RNA isolate was subjected to rRNA removal with the NEBNext rRNA Depletion Kit (NEB) using the manufacturer’s recommendations. The whole 10 µl of the RNA elute was used for reverse transcription with Superscript III (Invitrogen) using the recommended reaction supplemented with 0.5 μl of RiboLock RNase Inhibitor (Thermo Scientific) and 100 pmol of NNSR-RT primer with the following protocol: 45°C 30 min; 70°C, 15 min. The cDNA was bead-purified with 1.8 volume of SPRI Beads (Beckman Coulter), eluted in 27 μl of water and subjected to RNase-H (NEB) digestion at 37°C for 30 min followed by heat inactivation. After bead purification the 20 μl cDNA elute was used for 2nd strand synthesis in a 50 μl reaction containing: 1x NEB Buffer 2, 25 nmol dNTP, 5 Us of exo(-) Klenow Fragment (NEB), 200 pmol of NNSR-2 Primer for 30 min at 37°C. After bead purification half of the DNA elute was used for a 50-µl PCR reaction containing the NEBNext High-Fidelity 2x Master Mix (NEB), 25 pmol, each, of NNSR-Illumina and NNSR-nest-ind primers with the following cycling conditions: 98°C 10 sec; 5 cycles of 98°C 10 sec, 55°C 30 sec, 72°C 30 sec; 20 cycles of 98°C 10 sec, 65°C 30 sec, 72°C 30 sec; 72°C 5 min. 15 μl of the PCR reaction was separated on a 1% agarose gel and the smear of 500-700 bp was isolated. The indexed libraries were quantified by qPCR using the NEBNext Library Quant Kit for Illumia (NEB, mixed and sequenced on a MiSeq instrument (Illumina)) with a 2×250 paired-end setup.

### RNA sequencing analysis

Quality trimming and adapter removal were performed using fastp (v0.20.0 (42)). Read 1 and 2 adapter recognition sequences were provided for adapter removal (Illumina TruSeq Adapter Read 1: AGATCGGAAGAGCACACGTCTGAACTCCAGTCACNNNNNNATCTCGTATGCCGTCTTCTGCT TG, Illumina TruSeq Adapter Read 2: AGATCGGAAGAGCGTCGTGTAGGGAAAGAGTGT; NNNNNN: sample-specific index) and the leading two nucleotides were removed from each read (--trim_front1 2 --trim_front2 2). For quality trimming, bases in sliding windows with a mean quality below 30 (−5 -3 --cut_mean_quality 30) were discarded on both sides of the reads. Base correction in overlapping regions (-c) was applied. Reads with Ns and a length below-< 30 bp after trimming (-n 0 -l 30) were discarded.

Mapping was performed with BWA mem v 0.7.12-r1039 (43), using default parameters unless stated otherwise. Host-derived reads were removed by mapping quality controlled reads against the African green monkey genome (*Chlorocebus sabeus*, RefSeq assembly GCA_000409795.2), specifying the minimum seed length (-k 31). Unmapped reads were extracted using samtools v1.7 (44) and bamToFastq v2.17.0 (45), and subsequently mapped to the plasmid reference genomes of either MeV_vac2_-SARS2-S(H) or MeV_vac2_-SARS2-S(P), as appropriate. Host-free alignments were deduplicated using picard-tools MarkDuplicates (http://broadinstitute.github.io/picard) and left-aligned using GATK LeftAlignIndels v4.0 (46).

Sample majority consensus sequences were obtained by substituting minor frequency variants in the respective virus reference sequence for alternative variants with allele frequencies > 50%. Variant calling was performed with LoFreq v2.1.3 (47) using default parameters.

### Immunoperoxidase monolayer assay (IPMA)

For immunoperoxidase monolayer assay, Vero cells cultured in flat-bottom 12-well plates were fixed overnight with methanol at ™20°C two days after infection with a MOI of 0.01. The fixed cells were then washed three times with 1 mL PBS and subsequently blocked with PBS containing 2% bovine serum albumin (BSA) (Roth, Karlsruhe, Germany) for 30 min at 37°C. The cells were then probed for 1 h with a polyclonal rabbit anti-SARS-CoV-2-S protein antibody (1:2,250; ab252690; Abcam, Cambridge, UK) or a rabbit anti-MeV N protein antibody (1:1,000, ab23974, Abcam) in PBS with 2% BSA. The cells were washed 3 times with 1 ml PBS and subsequently incubated with the secondary HRP-coupled donkey anti-rabbit IgG(H+L) polyclonal antibody (1:1,000; 611-7202; Rockland, Gilbertsville, USA) for 1 h at 37°C. Then, the cells were washed 3 times, again. For detection, the cells were stained with TrueBlue peroxidase substrate solution (SeraCare, Milford, USA).

### Western Blot Analysis

Cells were lysed and immunoblotted as previously described (48). Rabbit anti-SARS-S protein antibody (1:3,000; ab252690; Abcam), rabbit anti-MeV-N protein polyclonal antibody (1:5,000; ab23974; Abcam), and a mouse anti-ß-actin antibody (1:5,000; ab6276; Abcam) were used. Donkey anti-rabbit IgG-HRP (H&L) polyclonal antibody (1:10,000; 611-7202; Rockland) and goat anti-mouse IgG-HRP (1:10,000; A2554-1ML; Merck, Darmstadt, Germany) served as secondary antibodies. Peroxidase activity was visualized with an enhanced chemiluminescence detection kit (Thermo Scientific, Bremen, Germany) on ChemiDoc MP Imaging System (Biorad, Dreieich, Germany).

### Animal experiments

All animal experiments were carried out in compliance with the regulations of German animal protection laws and as authorized by the RP Darmstadt in consideration of the ARRIVE guidelines. Six- to 12-week-old old, treatment-naive IFNAR^-/-^-CD46Ge mice (10) that are deficient for type I IFN receptor and transgenically express human CD46 were bred in-house under SPF conditions and regularly controlled by animal care takers and institutional veterinarians for general signs of well-being, and animal weight was additionally controlled once a week during the experiments. For the experiments, animals were randomized for age- and sex-matched groups and housed in IVC cages in groups of 3 to 5 animals with nist packs as environmental enrichment at room temperature with regular 12 h day and night intervalls. Group sizes were calculated based on statistical considerations to yield sufficient statistical power as authorized by the respective competent authority. These animals were inoculated intraperitoneally (i.p.) with 1×10^5^ TCID_50_ of recombinant vaccine viruses in 200 μl volume, or subcutaneously (s.c.) with 10 μg recombinant SARS-CoV-2 S protein (Sino Biological Europe, Eschborn, Germany) adjuvanted with 500 μg aluminium hydroxide (Alhydrogel adjuvant 2%, vac-alu-250, InvivoGen, San Diego, CA, USA) in 100 μl volume on days 0 and 28. 200 μl blood was collected on days 0, and 28, while final serum was collected on day 49 post initial immunization (p.i.). serum samples were stored at -20°C. Mice were euthanized on day 49 p.i., and splenocytes were harvested for assessment of cellular immune responses.

### Total IgG and IgG1-/IgG2a quantification

MeV bulk antigens (10 μg/mL; Virion Serion, Würzburg) or recombinant SARS-CoV-2 S protein (5 μg/mL) were coated in 50 µl carbonate buffer (Na_2_CO_3_ 30 mM; NaHCO_3_ 70 mM; pH 9.6) per well on Nunc Maxisorp® 96 well ELISA plates (ebioscience) and incubated overnight at 4°C. The plates were washed three times with 200 µl ELISA washing buffer (PBS, 0.1% Tween 20 (w/v)) and blocked with 100 μL Blocking buffer (PBS; 5% BSA; 0.1% Tween 20) for at least 2 h at room temperature. Mouse sera were 5-fold serially diluted in ELISA dilution buffer (PBS, 1% BSA, 0.1% Tween 20), and 50 μL/well were used for the assay. The plates were incubated at 37°C for 2 h and washed three times with ELISA washing buffer, followed by incubation with 50 µl/well of HRP-conjugated rabbit anti-mouse total IgG (1:1,000 in ELISA dilution buffer; P0260, Dako Agilent, Santa Clara, CA, USA), goat-anti-mouse IgG1 (1:8,000 in ELISA dilution buffer; ab97240, Abcam, Cambridge, UK), or goat-anti-mouse IgG2a (1:8,000 in ELISA dilution buffer; ab97245, Abcam) at room temperature for 1 h. Subsequently, the plates were washed four times and 100 μL TMB substrate (ebioscience) was added per well. The reaction was stopped by addition of 50 μL/well H_2_SO_4_ (1 N) and the absorbance at 450 nm (specific signal) and 630nm (reference wavelength) was measured.

### Th1/Th2 cytokine multiplex assay

Quantification of Th1/Th2 cytokines in supernatant of splenocytes was performed using mouse high sensitivity T cell magnetic bead panel assay (MHSTCMAG-70K, Merck, Darmstadt, Germany). 5×10^5^ isolated splenocytes were co-cultured with different stimuli in 200 μL RPMI + 10% FBS, 2 mM L-Gln, and 1% penicillin-streptomycin for 36 h. For re-stimulation of SARS-CoV-2 S protein-specific T cells, splenocytes were co-cultivated with 5×10^4^ DC2.4 dendritic cells, the corresponding cell line transgenically expressing SARS-CoV-2 S protein or medium alone. After 36 h, cells were spun down and supernatants were collected and stored at -20°C till assayed. For multiplex assay, cytokines were coupled over night to magnetic beads coated with capture antibodies, labeled with biotinylated detection antibody and incubated with Streptavidin-PE conjugate. Fluorescence was measured using MAGPIX with xPONENT software (Luminex Instruments, Thermo Scientific, Bremen, Germany).

### Neutralization Assay

Virus neutralizing titers (VNT) were quantified as described previously (7). Towards this, sera were serially diluted in 2-fold dilution steps in DMEM in duplicates. A total of 50 PFU of MV_vac2_-GFP(P) or 100 TCID_50_ of SARS-CoV-2 (isolate MUC-IMB1) were mixed with diluted sera and incubated at 37°C for 1 h. MeV or SARS-CoV-2 virus-serum suspensions were added to 1×10^4^ Vero or Vero E6 cells, respectively, seeded 4 h prior to the assay in 96-well plates and incubated for 4 days at 37°C. VNTs were calculated as the reciprocal of the highest mean dilution that abolished infection.

### IFN-γ ELISpot Analysis

Murine interferon gamma (IFN-γ) enzyme-linked immunosorbent spot (ELISpot) assays were performed using the Mouse IFN-γ ELISPOT Pair kit including capture and detection antibody (BD Bioscience, Franklin Lakes, NJ, USA) and HRP Streptavidin (BD Bioscience) for ELISpot detection in combination with multiscreen immunoprecipitation (IP) ELISpot polyvinylidene difluoride (PVDF) 96-well plates (Merck Millipore, Darmstadt, Germany) according to the manufacturer’s instructions. 5×10^5^ isolated splenocytes were co-cultured with different stimuli in 200 μL RPMI + 10% FBS, 2 mM L-Gln, and 1% penicillin-streptomycin for 36 h. For re-stimulation of SARS-CoV-2 S protein-specific T cells, splenocytes were co-cultivated with 5×10^4^ JAWSII, DC2.4 dendritic cells, or the corresponding cell lines transgenically expressing SARS-CoV-2 S protein. In parallel, splenocytes were stimulated with 10 μg/mL MeV bulk antigen (Virion Serion). For general T cell stimulation, 10 μg/mL concanavalin A (ConA, Sigma-Aldrich) was used, and as negative control, splenocytes were left untreated. After 36 h, cells were spun down, supernatants were removed, and cells were lysed in the wells by hypotonic shock. Plates were incubated with biotin-conjugated anti-IFN-γ detection antibodies and streptavidin-HRP according to the manufacturer’s instructions. 3-Amino-9-ethyl-carbazole (AEC; Sigma-Aldrich) was dissolved in N,N-dimethylformamide (Merck Millipore) and used for peroxidase-dependent staining. Spots were counted using an Eli.Scan ELISpot scanner (AE.L.VIS, Hamburg, Germany) and ELISpot analysis software Eli.Analyse V5.0 (AE.L.VIS).

### Intracellular cytokine staining

For flow cytometry-based analysis of cytokine expression by intracellular cytokine staining (ICS), splenocytes of vaccinated mice were isolated, and 2×10^6^ splenocytes per mouse were cultivated in 200 μL RPMI1640 + 10% FBS, 2 mM L-Gln, 1× non-essential amino acids (Biochrom), 10 mM HEPES, 1% penicillin-streptomycin, 50 μM β-mercaptoethanol, 10 μg/mL brefeldin A (Sigma-Aldrich) with DC2.4-SARS2-S cells as used for ELISpot analysis. For general T cell stimulation, 0.25 μg/mL tetradecanoylphorbol acetate (TPA, Sigma Aldrich) and 0.5 μg/mL ionomycin (Iono, Sigma-Aldrich) were used as positive control, and medium alone served as negative control. Splenocytes were stimulated for 5 h at 37°C. Subsequently, cells were stained with fixable viability dye eFluor450 (eBioscience), α-CD4-PE (1:2,000; Cat.-No. 553049 BD, Franklin Lakes, NJ, USA), α-CD8-FITC (1:500; Cat.-No. 553031, BD), and α-CD3-PerCPCy5.5 (1:500; Cat.-No. 550763, BD). Subsequent to permeabilization with Fixation/Permeabilization Solution (BD) and Perm/Wash Buffer (BD), cells were stained with α-IFN-γ-APC (1:500; Cat.-No. 554413, BD), α-IL-2-AlexaFluor700 (1:200; Cat.-No. 503818, Biolegend, San Diego, USA) and α-TNF-α-Pe-Cy7 (1:500; Cat.-No. 557644, BD). Cells were fixed with ice-cold 1% paraformaldehyde (PFA) in PBS and analyzed via flow cytometry using an LSRII SORP flow cytometer (BD) and DIVA software (BD).

### T cell proliferation assay

Splenocytes isolated three weeks after the second immunization were labeled with 0.5 μM carboxyfluorescein-succinimidyl-ester (CFSE) (ebioscience, Life Technologies, Carlsbad, CA, USA) as previously described (49). In brief, 5×10^5^ labelled cells were seeded in RPMI 1640 supplemented with 10% mouse serum, 2 mM L-Glutamine, 10 mM HEPES, 1% penicillin/streptomycin, and 100 μM 2-mercaptoethanol in 96-wells. 200 μL Medium containing 10 µg/ml Concanavalin A (Con A, Sigma-Aldrich), 10 μg/ml MeV bulk antigen (Virion Serion), or 5×10^3^ DC2.4-SARS2-S cells were added to each well, and cultured for 6 days. Medium and wild type DC2.4 and JAWSII cells served as controls. Stimulated cells were subsequently stained with CD3-PacBlue (1:50; clone 500A2; Invitrogen Life Technologies), CD8-APC (1:100; clone 53-6.7; ebioscience) and CD4-PE (1:2000; Cat. 553049; BD) antibodies and fixed with 1% PFA in PBS. Finally, the stained cells were analyzed by flow cytometry using an LSR II flow cytometer (BD) and FCS Express software (De Novo Software).

### CTL killing assay

For re-stimulation of T cells isolated 3 weeks after the second immunization, 5×10^6^ splenocytes were co-cultured with 5×10^4^ DC2.4-SARS2-S cells for 6 days in 12-wells in RPMI 1640 supplemented with 10% FBS, 2 nM L-Glutamin, 1 mM HEPES, 1% penicillin/streptomycin, 100 μM 2-mercaptoethanol, and 100 U/ml murine rIL-2 (Peprotech, Hamburg, Germany). 5×10^3^ EL-4_red_ cells were labeled with 0.5 µM CFSE and mixed with 5×10^3^ EL-4_green_-SARS2-S cells per well. Splenocytes were counted and co-cultured with EL-4 target cells at the indicated ratios for 4 h at 37°C. Afterwards, EL-4 cells were labeled with Fixable Viability Dye eFluor® 780 (ebioscience), fixed with 1% paraformaldehyde (PFA), and analyzed by flow cytometry using an LSR II flow cytometer (BD) and FCS Express. For indication of Antigen:NC EL-4 ratio the cell count of viable SARS-CoV-2 S-expressing cells was divided by the population of viable negative controls.

## Statistical analyses

To compare the means of different groups in growth kinetics, a non-parametric One-way ANOVA was performed. For ICS analysis, the non-parametric two-tailed Mann-Whitney test was used to compare cytokines levels between DC2.4 and DC2.4-SARS2-S-restimulated splenoycytes within the MeV-_vac2_-SARS-2-S(H) vaccine group. Note, that these exploratory analyses have been done without correction for multiple testing. For proliferation assay the mean differences were calculated and analyzed using one-tailed Mann-Whitney t-test. To all three groups in CTL killing assays a linear curve was fitted for antigen vs. logarithmised effector-target ratio E:T. The p values testing for differences in slopes were calculated and populations of SARS2-S(H) compared with control ATU vaccinated cells. The p values were not adjusted for multiplicity due to the explorative character of the study. For VNT and fusion activity statistical analysis, one-way ANOVA was performed in combination with Tukey’s Multi comparison test to compare all pair means. For multiplex statistical analysis, two-way ANOVA analysis was applied with paired Tukey’s Multi comparison test as post hoc test. For statistical analysis of grouped ELISpot data, two-way ANOVA analysis was applied with paired Tukey’s Multi comparison test.

## Acknowledgments

This work was supported by the German Center for Infection Research (DZIF; TTU 01.805, TTU 01.922_00). The authors would like to thank Daniela Müller and Carina Kruip for excellent technical assistance, Björn Becker for assistance in multiplex analysis, Csabas Miskey for assistance with NGS, Christel Kamp for excellent advice on statistics, and Marcel Rommel for cell sorting. The authors are indebted to Gerhard Dobler for providing SARS-CoV-2 isolate MUC-IMB1, Maria Vehreschild for human patient reconvalescent serum, Kenneth Rock for DC2.4 cells, Roberto Cattaneo for providing the pBR(+)MV_vac2_ construct, and Urs Schneider for providing the PolII rescue system used to generate and to rescue recombinant MeV vectors. The authors would further like to thank Bakhos Tannous for providing pCSCW2gluc-IRES-GFP. Moreover, the authors would like to thank Roberto Cattaneo and Veronika von Messling for valuable comments on the manuscript.

## Supplementary Figures

**Suppl. Fig. S1:**
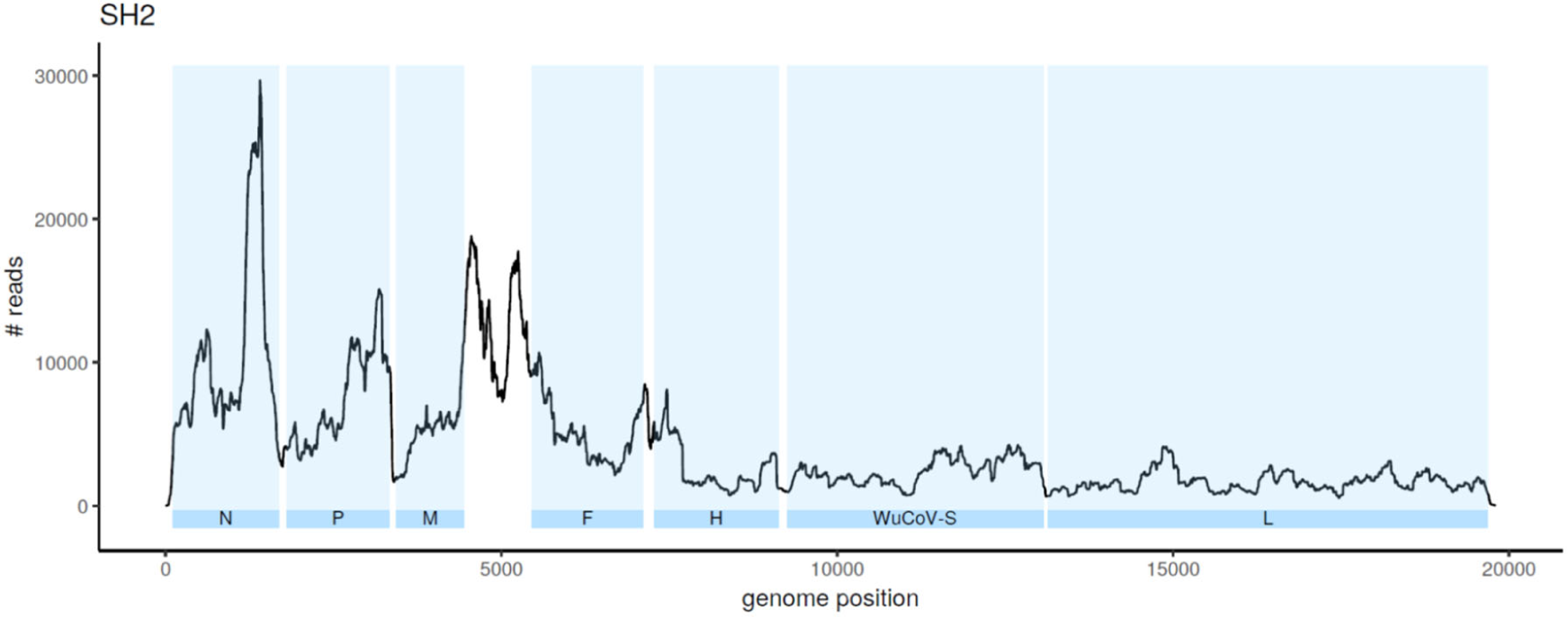
Coverage of of vaccine candidate MeV_vac2_-SARS2-S(H) genome during next generation sequencing. Schematic depiction of read frequency at each position of the vaccine virus genome. Blue areas indicate respective viral coding sequences, white areas indicate intergenic regions and untranscribed terminal regions (UTRs) of the genome. Coverage across the genome was sufficient for variant detection and reflects the transcription gradient typically observed in measles virus total RNAseq data. Since the majority of reads are mRNA-derived, low read numbers decrease strongly between the coding regions and continually towards the 5’ end.

**Suppl. Fig. S2:**
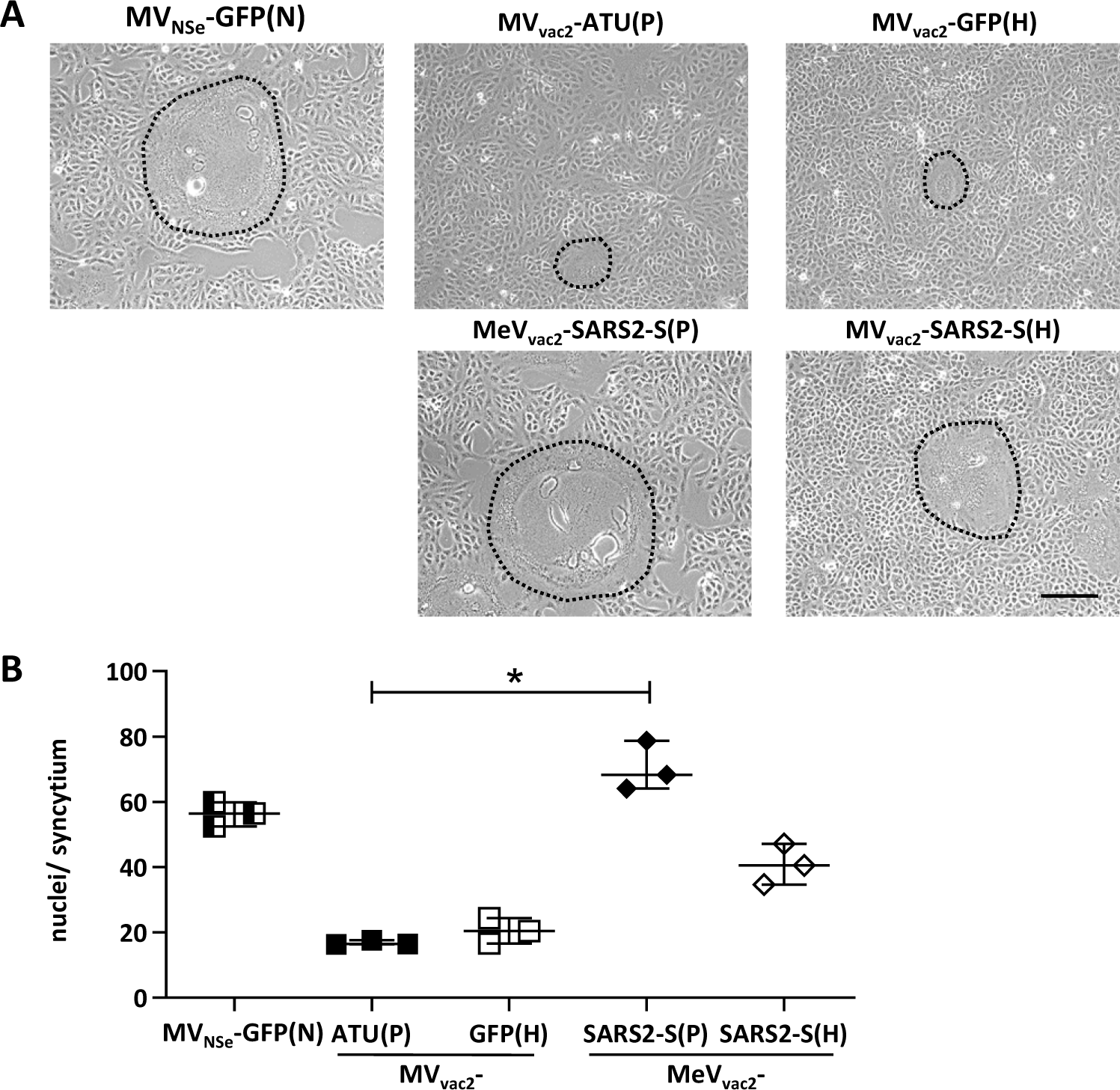
Characterization of fusogenic phenotype of MeV_vac2_-SARS2-S(P) and MeV_vac2_-SARS2-S(H). **(A**) Photographs of fusion activity of Vero cells infected at an MOI of 0.01 with MeV_vac2_-SARS2-S(P) or MeVvac2-SARS2-S(H) encoding SARS-CoV-2 S in additional transcription units post P or post H, respectively, in direct comparison to MV_vac2_-ATU(P) or MV_vac2_-GFP(H) control vaccine viruses or MV_NSe_-GFP(N) hyperfusogenic oncolytic MeV. Representative picture of one out of three independent experiments. Scale bar represents 200 mm. (**B**) Cell fusion was quantified 30 h after infection. For statistical analysis, one-way ANOVA was performed in combination with Tukey’s Multi comparison test to compare all pair means. *, p<0.05.

**Suppl. Fig. S3:**
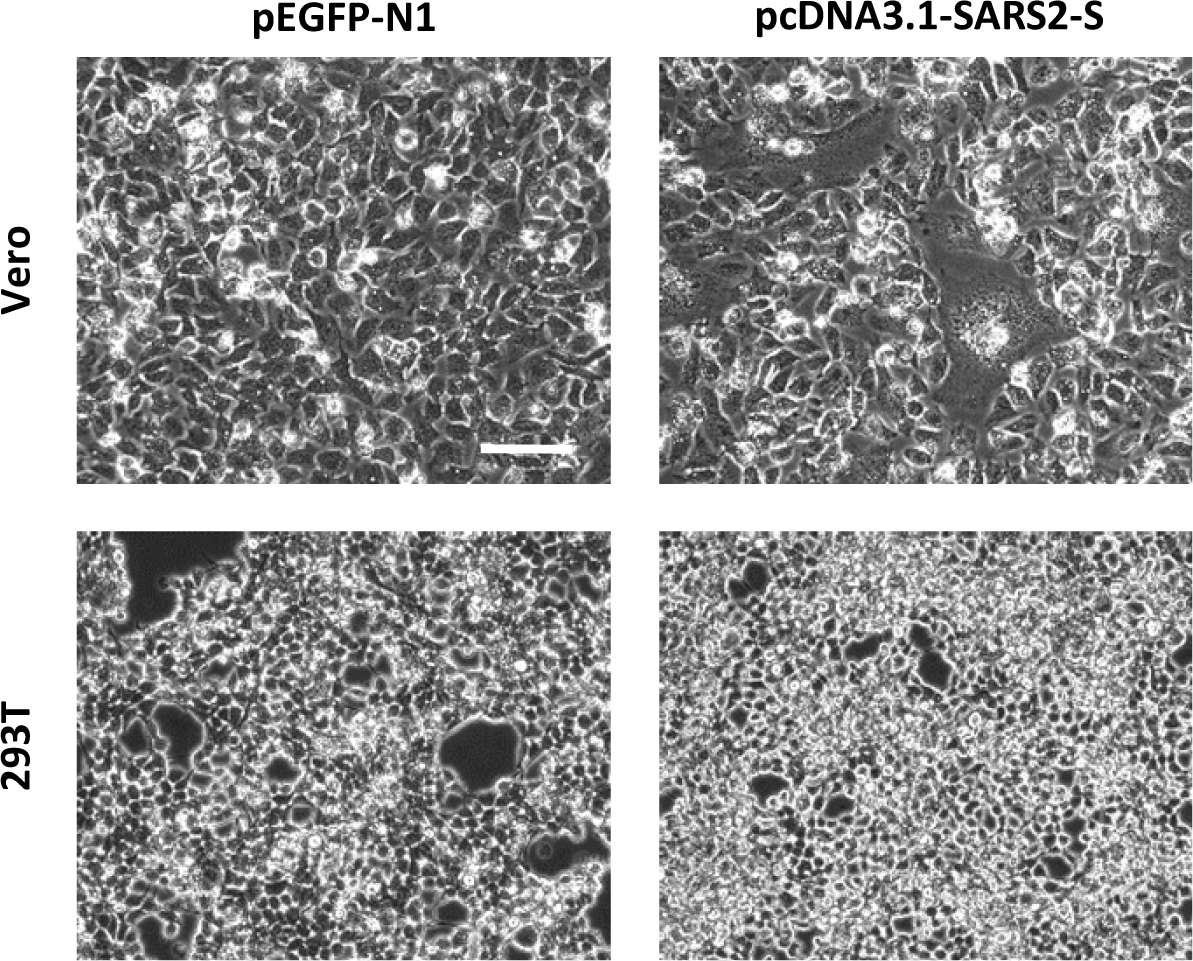
Expression of SARS-CoV-2 S protein in Vero and 293T cells. Photographic depiction of fusion activity in Vero or 293T cells 48 h after transfection with 1 µg of SARS-CoV-2 S expression plasmid of control DNA. One representative out of three independent experiments is shown. Scale bar represents 100 mm.

**Suppl. Fig. S4:**
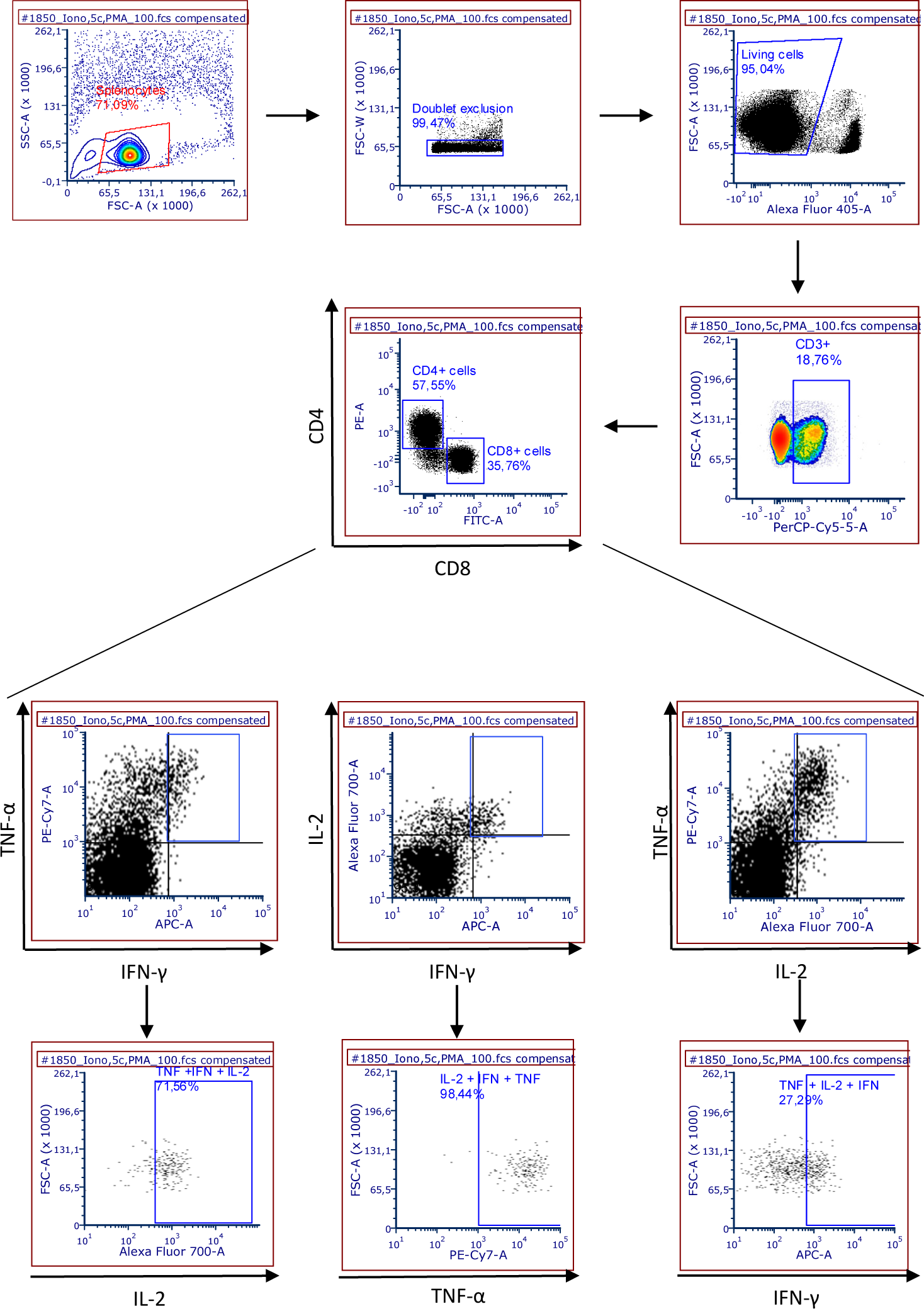
Gating strategy for intracellular cytokine staining. Exemplary depiction of the gating strategy to analyze T cells after re-stimulation and staining for cytokine induction. The gating strategy includes cell doublet exclusion, selection for living cells and separation of CD8+ and CD4+ T cells within CD3+ splenocyte populations. Respectively gated T cell populations were then analysed for expression of IFN-g, TNF-a, or IL-2. Multi-colour flow cytometry allows assessment of double- or triple-positive cells, exemplarily shown for CD4+ T cells after stimulation with ionomycin and PMA.

## References

1. F. Wu, S. Zhao, B. Yu, Y.-M. Chen, W. Wang, Z.-G. Song, Y. Hu, Z.-W. Tao, J.-H. Tian, Y.-Y. Pei, M.-L. Yuan, Y.-L. Zhang, F.-H. Dai, Y. Liu, Q.-M. Wang, J.-J. Zheng, L. Xu, E. C. Holmes, Y.-Z. Zhang, A new coronavirus associated with human respiratory disease in China. Nature 579, 265–269 (2020).

2. Q. Li, X. Guan, P. Wu, X. Wang, L. Zhou, Y. Tong, R. Ren, K. S. M. Leung, E. H. Y. Lau, J. Y. Wong, X. Xing, N. Xiang, Y. Wu, C. Li, Q. Chen, D. Li, T. Liu, J. Zhao, M. Liu, W. Tu, C. Chen, L. Jin, R. Yang, Q. Wang, S. Zhou, R. Wang, H. Liu, Y. Luo, Y. Liu, G. Shao, H. Li, Z. Tao, Y. Yang, Z. Deng, B. Liu, Z. Ma, Y. Zhang, G. Shi, T. T. Y. Lam, J. T. Wu, G. F. Gao, B. J. Cowling, B. Yang, G. M. Leung, Z. Feng, Early Transmission Dynamics in Wuhan, China, of Novel Coronavirus-Infected Pneumonia. The New England journal of medicine 382, 1199–1207 (2020).

3. R. Li, S. Pei, B. Chen, Y. Song, T. Zhang, W. Yang, J. Shaman, Substantial undocumented infection facilitates the rapid dissemination of novel coronavirus (SARS-CoV-2). 10.5281/ZENODO.3699624 (2020).

4. M. D. Mühlebach, Vaccine platform recombinant measles virus. Virus genes 53, 733–740 (2017).

5. N. Escriou, B. Callendret, V. Lorin, C. Combredet, P. Marianneau, M. Février, F. Tangy, Protection from SARS coronavirus conferred by live measles vaccine expressing the spike glycoprotein. Virology 452-453, 32–41 (2014).

6. M. Liniger, A. Zuniga, A. Tamin, T. N. Azzouz-Morin, M. Knuchel, R. R. Marty, M. Wiegand, S. Weibel, D. Kelvin, P. A. Rota, H. Y. Naim, Induction of neutralising antibodies and cellular immune responses against SARS coronavirus by recombinant measles viruses. Vaccine 26, 2164–2174 (2008).

7. A. H. Malczyk, A. Kupke, S. Prüfer, V. A. Scheuplein, S. Hutzler, D. Kreuz, T. Beissert, S. Bauer, S. Hubich-Rau, C. Tondera, H. S. Eldin, J. Schmidt, J. Vergara-Alert, Y. Süzer, J. Seifried, K.-M. Hanschmann, U. Kalinke, S. Herold, U. Sahin, K. Cichutek, Z. Waibler, M. Eickmann, S. Becker, M. D. Mühlebach, A Highly Immunogenic and Protective Middle East Respiratory Syndrome Coronavirus Vaccine Based on a Recombinant Measles Virus Vaccine Platform. Journal of virology 89, 11654–11667 (2015).

8. B. S. Bodmer, A. H. Fiedler, J. R. H. Hanauer, S. Prüfer, M. D. Mühlebach, Live-attenuated bivalent measles virus-derived vaccines targeting Middle East respiratory syndrome coronavirus induce robust and multifunctional T cell responses against both viruses in an appropriate mouse model. Virology 521, 99–107 (2018).

9. S. Heidmeier, J. R. H. Hanauer, K. Friedrich, S. Prüfer, I. C. Schneider, C. J. Buchholz, K. Cichutek, M. D. Mühlebach, A single amino acid substitution in the measles virus F2 protein reciprocally modulates membrane fusion activity in pathogenic and oncolytic strains (2014).

10. B. Mrkic, J. Pavlovic, T. Rülicke, P. Volpe, C. J. Buchholz, D. Hourcade, J. P. Atkinson, A. Aguzzi, R. Cattaneo, Measles Virus Spread and Pathogenesis in Genetically Modified Mice (1998).

11. P. Marrack, A. S. McKee, M. W. Munks, Towards an understanding of the adjuvant action of aluminium. Nature reviews. Immunology 9, 287–293 (2009).

12. P. He, Y. Zou, Z. Hu, Advances in aluminum hydroxide-based adjuvant research and its mechanism. Human vaccines & immunotherapeutics 11, 477–488 (2015).

13. R.-D. Jiang, M.-Q. Liu, Y. Chen, C. Shan, Y.-W. Zhou, X.-R. Shen, Q. Li, L. Zhang, Y. Zhu, H.-R. Si, Q. Wang, J. Min, X. Wang, W. Zhang, B. Li, H.-J. Zhang, R. S. Baric, P. Zhou, X.-L. Yang, Z.-L. Shi, Pathogenesis of SARS-CoV-2 in Transgenic Mice Expressing Human Angiotensin-Converting Enzyme 2. Cell. 10.1016/j.cell.2020.05.027 (2020).

14. M. Mura, C. Ruffié, E. Billon-Denis, C. Combredet, J. N. Tournier, F. Tangy, hCD46 receptor is not required for measles vaccine Schwarz strain replication in vivo: Type-I IFN is the species barrier in mice. Virology 524, 151–159 (2018).

15. A. Le Bon, G. Schiavoni, G. D’Agostino, I. Gresser, F. Belardelli, D. F. Tough, Type I Interferons Potently Enhance Humoral Immunity and Can Promote Isotype Switching by Stimulating Dendritic Cells In Vivo (2001).

16. I. J. Amanna, N. E. Carlson, M. K. Slifka, Duration of Humoral Immunity to Common Viral and Vaccine Antigens (2007).

17. S. Carryn, M. Feyssaguet, M. Povey, E. Di Paolo, Long-term immunogenicity of measles, mumps and rubella-containing vaccines in healthy young children: A 10-year follow-up. Vaccine 37, 5323–5331 (2019).

18. S. J. Russell, M. J. Federspiel, K.-W. Peng, C. Tong, D. Dingli, W. G. Morice, V. Lowe, M. K. O’Connor, R. A. Kyle, N. Leung, F. K. Buadi, S. V. Rajkumar, M. A. Gertz, M. Q. Lacy, A. Dispenzieri, Remission of disseminated cancer after systemic oncolytic virotherapy. Mayo Clinic proceedings 89, 926–933 (2014).

19. P. Msaouel, M. Opyrchal, A. Dispenzieri, K. W. Peng, M. J. Federspiel, S. J. Russell, E. Galanis, Clinical Trials with Oncolytic Measles Virus: Current Status and Future Prospects. Current cancer drug targets 18, 177–187 (2018).

20. A. Dispenzieri, C. Tong, B. LaPlant, M. Q. Lacy, K. Laumann, D. Dingli, Y. Zhou, M. J. Federspiel, M. A. Gertz, S. Hayman, F. Buadi, M. O’Connor, V. J. Lowe, K.-W. Peng, S. J. Russell, Phase I trial of systemic administration of Edmonston strain of measles virus genetically engineered to express the sodium iodide symporter in patients with recurrent or refractory multiple myeloma. Leukemia 31, 2791–2798 (2017).

21. D. E. Griffin, W.-H. W. Lin, A. N. Nelson, Understanding the causes and consequences of measles virus persistence (2018).

22. K. Ramsauer, M. Schwameis, C. Firbas, M. Müllner, R. J. Putnak, S. J. Thomas, P. Desprès, E. Tauber, B. Jilma, F. Tangy, Immunogenicity, safety, and tolerability of a recombinant measles-virus-based chikungunya vaccine: a randomised, double-blind, placebo-controlled, active-comparator, first-in-man trial. The Lancet Infectious Diseases 15, 519–527 (2015).

23. E. C. Reisinger, R. Tschismarov, E. Beubler, U. Wiedermann, C. Firbas, M. Loebermann, A. Pfeiffner, M. Muellner, E. Tauber, K. Ramsauer, Immunogenicity, Safety, and Tolerability of the Measles-Vectored Chikungunya Virus Vaccine MV-CHIK: A Double-Blind, Randomised, Placebo-Controlled and Active-Controlled Phase 2 Trial (2019).

24. M. Hoffmann, H. Kleine-Weber, S. Schroeder, N. Krüger, T. Herrler, S. Erichsen, T. S. Schiergens, G. Herrler, N.-H. Wu, A. Nitsche, M. A. Müller, C. Drosten, S. Pöhlmann, SARS-CoV-2 Cell Entry Depends on ACE2 and TMPRSS2 and Is Blocked by a Clinically Proven Protease Inhibitor. Cell 181, 271-280.e8 (2020).

25. D. Wrapp, Wang, Nianshuang, Corbett, Kizzmekia S., J. A. Goldsmith, C.-L. Hsieh, O. Abiona, B. S. Graham, J. S. McLellan, Cryo-EM structure of the 2019-nCoV spike in the prefusion conformation 60 (2020).

26. J. Yu, L. H. Tostanoski, L. Peter, N. B. Mercado, K. McMahan, S. H. Mahrokhian, J. P. Nkolola, J. Liu, Z. Li, A. Chandrashekar, D. R. Martinez, C. Loos, C. Atyeo, S. Fischinger, J. S. Burke, M. D. Slein, Y. Chen, A. Zuiani, F. J. N Lelis, M. Travers, S. Habibi, L. Pessaint, A. van Ry, K. Blade, R. Brown, A. Cook, B. Finneyfrock, A. Dodson, E. Teow, J. Velasco, R. Zahn, F. Wegmann, E. A. Bondzie, G. Dagotto, M. S. Gebre, X. He, C. Jacob-Dolan, M. Kirilova, N. Kordana, Z. Lin, L. F. Maxfield, F. Nampanya, R. Nityanandam, J. D. Ventura, H. Wan, Y. Cai, B. Chen, A. G. Schmidt, D. R. Wesemann, R. S. Baric, G. Alter, H. Andersen, M. G. Lewis, D. H. Barouch, DNA vaccine protection against SARS-CoV-2 in rhesus macaques. Science (New York, N.Y.). 10.1126/science.abc6284 (2020).

27. N. M. A. Okba, M. A. Müller, W. Li, C. Wang, C. H. GeurtsvanKessel, V. M. Corman, M. M. Lamers, R. S. Sikkema, E. de Bruin, F. D. Chandler, Y. Yazdanpanah, Q. Le Hingrat, D. Descamps, N. Houhou-Fidouh, C. B. E. M. Reusken, B.-J. Bosch, C. Drosten, M. P. G. Koopmans, B. L. Haagmans, Severe Acute Respiratory Syndrome Coronavirus 2-Specific Antibody Responses in Coronavirus Disease Patients. Emerging infectious diseases 26, 1478–1488 (2020).

28. F. A. Rey, K. Stiasny, M.-C. Vaney, M. Dellarole, F. X. Heinz, The bright and the dark side of human antibody responses to flaviviruses: lessons for vaccine design. EMBO reports 19, 206–224 (2018).

29. L. Liu, Q. Wei, Q. Lin, J. Fang, H. Wang, H. Kwok, H. Tang, K. Nishiura, J. Peng, Z. Tan, T. Wu, K.-W. Cheung, K.-H. Chan, X. Alvarez, C. Qin, A. Lackner, S. Perlman, K.-Y. Yuen, Z. Chen, Anti-spike IgG causes severe acute lung injury by skewing macrophage responses during acute SARS-CoV infection. JCI insight 4 (2019).

30. D. Naniche, Human immunology of measles virus infection. Curr Top Microbiol Immunol. (2009).

31. L. Bao, W. Deng, B. Huang, H. Gao, J. Liu, L. Ren, Q. Wei, P. Yu, Y. Xu, F. Qi, Y. Qu, F. Li, Q. Lv, W. Wang, J. Xue, S. Gong, M. Liu, G. Wang, S. Wang, Z. Song, L. Zhao, P. Liu, L. Zhao, F. Ye, H. Wang, W. Zhou, N. Zhu, W. Zhen, H. Yu, X. Zhang, L. Guo, L. Chen, C. Wang, Y. Wang, X. Wang, Y. Xiao, Q. Sun, H. Liu, F. Zhu, C. Ma, L. Yan, M. Yang, J. Han, W. Xu, W. Tan, X. Peng, Q. Jin, G. Wu, C. Qin, The pathogenicity of SARS-CoV-2 in hACE2 transgenic mice. Nature. 10.1038/s41586-020-2312-y (2020).

32. S. F. Sia, L.-M. Yan, A. W. H. Chin, K. Fung, K.-T. Choy, A. Y. L. Wong, P. Kaewpreedee, R. A. P. M. Perera, L. L. M. Poon, J. M. Nicholls, M. Peiris, H.-L. Yen, Pathogenesis and transmission of SARS-CoV-2 in golden hamsters. Nature. 10.1038/s41586-020-2342-5 (2020).

33. J. F.-W. Chan, A. J. Zhang, S. Yuan, V. K.-M. Poon, C. C.-S. Chan, A. C.-Y. Lee, W.-M. Chan, Z. Fan, H.-W. Tsoi, L. Wen, R. Liang, J. Cao, Y. Chen, K. Tang, C. Luo, J.-P. Cai, K.-H. Kok, H. Chu, K.-H. Chan, S. Sridhar, Z. Chen, H. Chen, K. K.-W. To, K.-Y. Yuen, Simulation of the clinical and pathological manifestations of Coronavirus Disease 2019 (COVID-19) in golden Syrian hamster model: implications for disease pathogenesis and transmissibility (2020).

34. Z. Shen, G. Reznikoff, G. Dranoff, K. L. Rock, Cloned dendritic cells can present exogenous antigens on both MHC class I and class II molecules. Journal of immunology (Baltimore, Md. : 1950) 158, 2723–2730 (1997).

35. K. Friedrich, J. R. Hanauer, S. Prüfer, R. C. Münch, I. Völker, C. Filippis, C. Jost, K.-M. Hanschmann, R. Cattaneo, K.-W. Peng, A. Plückthun, C. J. Buchholz, K. Cichutek, M. D. Mühlebach, DARPin-targeting of measles virus: unique bispecificity, effective oncolysis, and enhanced safety. Molecular therapy : the journal of the American Society of Gene Therapy 21, 849–859 (2013).

36. J. W. Hewett, B. Tannous, B. P. Niland, F. C. Nery, J. Zeng, Y. Li, X. O. Breakefield, Mutant torsinA interferes with protein processing through the secretory pathway in DYT1 dystonia cells. Proceedings of the National Academy of Sciences of the United States of America 104, 7271–7276 (2007).

37. R. Zufferey, D. Nagy, R. J. Mandel, L. Naldini, D. Trono, Multiply attenuated lentiviral vector achieves efficient gene delivery in vivo (1997).

38. R. C. Münch, M. D. Mühlebach, T. Schaser, S. Kneissl, C. Jost, A. Plückthun, K. Cichutek, C. J. Buchholz, DARPins: an efficient targeting domain for lentiviral vectors. Molecular therapy : the journal of the American Society of Gene Therapy 19, 686–693 (2011).

39. A. Martin, P. Staeheli, U. Schneider, RNA polymerase II-controlled expression of antigenomic RNA enhances the rescue efficacies of two different members of the Mononegavirales independently of the site of viral genome replication. Journal of virology 80, 5708–5715 (2006).

40. G. Kaerber, Beitrag zur kollektiven Behandlung pharmakologischer Reihenversuche (1931).

41. J. R. del Valle, P. Devaux, G. Hodge, N. J. Wegner, M. B. McChesney, R. Cattaneo, A vectored measles virus induces hepatitis B surface antigen antibodies while protecting macaques against measles virus challenge. Journal of virology 81, 10597–10605 (2007).

42. S. Chen, Y. Zhou, Y. Chen, J. Gu, fastp: an ultra-fast all-in-one FASTQ preprocessor. Bioinformatics (Oxford, England) 34, i884–i890 (2018).

43. H. Li, R. Durbin, Fast and accurate short read alignment with Burrows-Wheeler transform. Bioinformatics (Oxford, England) 25, 1754–1760 (2009).

44. H. Li, B. Handsaker, A. Wysoker, T. Fennell, J. Ruan, N. Homer, G. Marth, G. Abecasis, R. Durbin, The Sequence Alignment/Map format and SAMtools. Bioinformatics (Oxford, England) 25, 2078–2079 (2009).

45. A. R. Quinlan, I. M. Hall, BEDTools: a flexible suite of utilities for comparing genomic features. Bioinformatics (Oxford, England) 26, 841–842 (2010).

46. A. McKenna, M. Hanna, E. Banks, A. Sivachenko, K. Cibulskis, A. Kernytsky, K. Garimella, D. Altshuler, S. Gabriel, M. Daly, M. A. DePristo, The Genome Analysis Toolkit: a MapReduce framework for analyzing next-generation DNA sequencing data. Genome research 20, 1297–1303 (2010).

47. A. Wilm, P. P. K. Aw, D. Bertrand, G. H. T. Yeo, S. H. Ong, C. H. Wong, C. C. Khor, R. Petric, M. L. Hibberd, N. Nagarajan, LoFreq: a sequence-quality aware, ultra-sensitive variant caller for uncovering cell-population heterogeneity from high-throughput sequencing datasets. Nucleic acids research 40, 11189–11201 (2012).

48. S. Funke, A. Maisner, M. D. Mühlebach, U. Koehl, M. Grez, R. Cattaneo, K. Cichutek, C. J. Buchholz, Targeted cell entry of lentiviral vectors. Molecular therapy : the journal of the American Society of Gene Therapy 16, 1427–1436 (2008).

49. A. B. Lyons, C. R. Parish, Determination of lymphocyte division by flow cytometry. Journal of immunological methods 171, 131–137 (1994).

